# Initiation of a ZAKα-dependent Ribotoxic Stress Response by the Innate Immunity Endoribonuclease RNase L

**DOI:** 10.1101/2023.10.12.562082

**Authors:** Jiajia Xi, Goda Snieckute, Abhishek Asthana, Christina Gaughan, Simon Bekker-Jensen, Robert H. Silverman

## Abstract

RNase L is a regulated endoribonuclease in higher vertebrates that functions in antiviral innate immunity. Interferons induce OAS enzymes that sense double-stranded RNA of viral origin leading to synthesis of 2’,5’-oligoadenylate (2-5A) activators of RNase L. However, it is unknown precisely how RNase L inhibits viral infections. To isolate effects of RNase L from other effects of double-stranded RNA or virus, 2-5A was directly introduced into cells. Here we report that RNase L activation by 2-5A causes a ribotoxic stress response that requires the ribosome-associated MAP3K, ZAKα. Subsequently, the stress-activated protein kinases (SAPK) JNK and p38α are phosphorylated. RNase L activation profoundly altered the transcriptome by widespread depletion of mRNAs associated with different cellular functions, but also by SAPK-dependent induction of inflammatory genes. Our findings show that 2-5A is a ribotoxic stressor that causes RNA damage through RNase L triggering a ZAKα kinase cascade leading to proinflammatory signaling and apoptosis.

**Highlights:**

- RNase L signals antiviral innate immunity from the ribosome
- The ribotoxic stress response requires the MAP3K ZAKα
- RNA cleavage leads to transcription of proinflammatory genes
- ZAKα contributes to apoptosis downstream of RNase L activity

## INTRODUCTION

A wide range of different cellular insults trigger stress responses that determine cell survival and cell fate ^1^. Adaptation and survival of the injured cell is accompanied by inflammatory signaling. However, the cell dies when the damage exceeds the capacity of the cell to repair. A ribotoxic stress response is a type of cellular stress pathway in which RNA damage or translational insults cause ribosomes to stall and collide ^1–4^. Ribotoxic stressors include ribotoxins that cleave 28S rRNA (Shiga toxin, α-sarcin and ricin), some, but not all, antibiotics that inhibit translational elongation (anisomycin, cycloheximide), and ultraviolet radiation (uv) that damages mRNA^1,4,5^. Ribotoxic stress is sensed by the MAP3K ZAKα, a mixed-lineage kinase that interacts with actively translating ribosomes. Following activation, ZAKα phosphorylates the MAP2Ks 3/6, 4/7 ^6^. Subsequently, the MAP2Ks phosphorylate the stress-activated MAPKs (SAPKs), JNK and p38α. There are two ZAK isoforms, a longer form, ZAKα, with a flexible C-terminal segment that contains two regions which binds to ribosomes, and a short form, ZAKβ that neither associates with ribosomes nor is involved in the ribotoxic stress response ^3^. ZAKα phosphorylation in response to ribotoxic stress leads to its dissociation from ribosomes and phosphorylation of SAPKs ^3^. Activation of JNK and p38α stimulates proinflammatory signaling, while activation of JNK also leads to apoptosis ^7^.

RNase L is a regulated endoribonuclease that functions in antiviral innate immunity downstream of the interferon (IFN)-induced 2’,5’-oligoadenylate synthetases (OASs) ^8,9^. Double-stranded RNA (dsRNA), of either cellular or viral origin, activate IFN-inducible OASs1-3 ^10,11^. Once activated by dsRNA, these OASs produce 5’-phosphorylated 2’,5’-oligoadenylates (known as 2-5A) from ATP ^9,12,13^. 2-5A binding to RNase L in the presence of ATP causes its dimerization and activation ^14–16^. The antiviral activity of RNase L is accompanied by degradation of viral and cellular single-strand RNAs (ssRNAs) ^17^. RNase L is a general endoribonuclease that cleaves ssRNAs predominantly at UpUp^N and UpAp^N sites, restricting protein synthesis through endonucleolytic cleavage of actively translated mRNAs ^18–23^. Moreover, RNase L also cleaves rRNA in intact ribosomes, as well as some tRNAs ^20,24,25^. Interestingly, RNase L mediated cleavage of cellular or viral ssRNAs leads to phosphorylation of SAPKs JNK and p38α ^22,26^. In addition, JNK activation by RNase L beyond a poorly defined threshold leads to apoptosis ^22,27^. Here we show that 2-5A activation of RNase L leads to imbalanced transcriptome changes and a ZAKα-dependent ribotoxic stress response involving proinflammatory signaling and apoptosis.

## RESULTS

### The MAP3K ZAKα is required for the 2-5A/RNase L-induced ribotoxic stress response

To specifically activate RNase L, enzymatically-synthesized and HPLC-purified trimer 2-5A was transfected into WT mouse bone marrow macrophages (BMMs) (Figs. 1 and S1). 2-5A stimulated RNase L in a dose and time dependent manner (≥ 2 μM; ≥2 h) as determined by appearance of characteristic, discrete rRNA cleavage products (Fig. 1A,B) ^24,25^. Breakdown of 28S and 18S rRNA was observed in WT BMM transfected with 2-5A, but not with dephosphorylated 2-5A (A3), included as a negative control, whereas neither 2-5A or A3 induced rRNA cleavage in RNase L KO BMM (Fig. 1C). Therefore, rRNA cleavage induced by 2-5A was due to RNase L activity, and only in response to authentic 2-5A. In addition, prolonged incubation with 2-5A (> 5h) led to death of WT BMMs, but not of RNase L KO BMMs (Fig. 1D).

**Figure 1.**
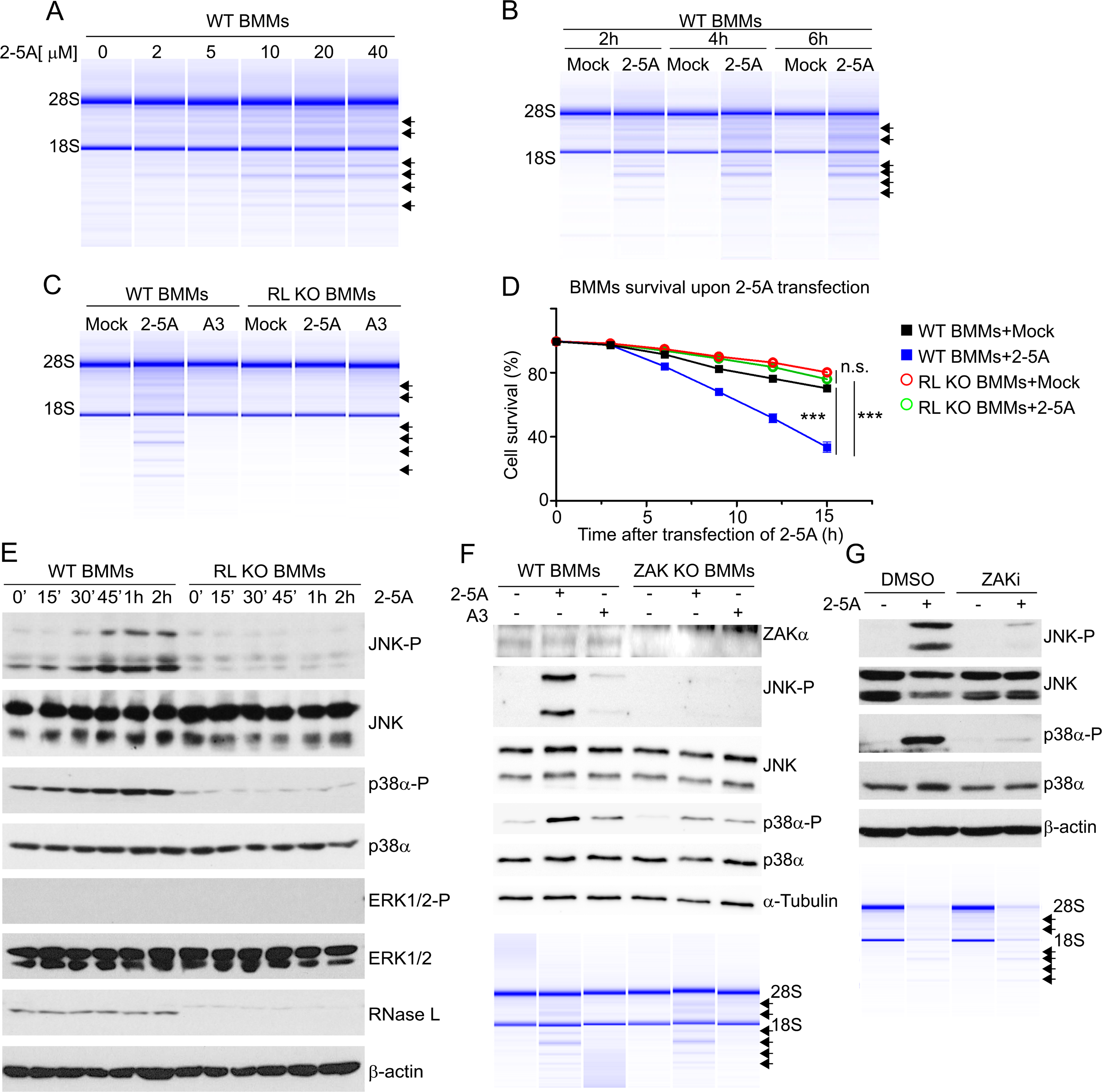
Activation of RNase L by 2-5A leads to ZAKα-dependent SAPK phosphorylation and cell death. (A) Dose- and (B) time-dependent induction of RNase L activity by 2-5A in WT BMMs as determined by rRNA cleavages. (C) RNase L mediated rRNA cleavage in WT BMM, but not in RNase L KO BMMs, by transfection with 2-5A (20 μM) for 3 h in comparison to mock- or A3-transfection controls. (D) Cell survival as determined by real-time imaging of stained WT or RNase L KO BMMs that were either mock transfected or transfected with 2-5A (20 μM). (E) Total and phosphorylated JNK, p38α, and ERK 1/2, and RNase L and β-actin in 2-5A-transfected WT BMM and RNase L KO BMM as determined by Western blotting. (F) Levels of ZAKα, total and phosphorylated JNK and p38α, and α-tubulin in WT BMM and ZAK KO BMM transfected with 2-5A (20 μM) or A3 (20 μM) for 2 h as determined by Western blotting. Lower panel shows intact and cleaved rRNA as determined in an RNA chip (Agilent). (G) Levels of total and phosphorylated JNK and p38α and β-actin in WT BMM treated with DMSO or ZAK inhibitior (ZAKi) compound 3h in either mock-transfected or 2-5A-transfected (20 μM) WT BMM for 2 h as determined by Western blotting. Lower panel, intact and cleaved rRNA in an RNA chip (Agilent).

Previously, we reported RNase L activation induced JNK phosphorylation in response to dsRNA, 2-5A or virus-infection ^22,26^. To probe the molecular mechanism of the 2-5A/RNase L-induced ribotoxic stress response, phosphorylation of JNK, p38α and ERK1/2 was monitored during 2-5A transfections. In WT BMM, but not in RNase L KO BMM, 2-5A induced phosphorylation of JNK beginning at 30-45 min (Fig. 1E). There were also low basal levels of p38α phosphorylation in the WT BMM which were increased following 2-5A transfection within a similar time frame. However, ERK1/2 were not phosphorylated in response to 2-5A transfection of WT BMM, suggesting that the 2-5A/RNase L-induced ribotoxic stress caused JNK- and p38α-, but not ERK-signaling.

Because the ribosome-associated MAP3K ZAKα phosphorylates MAP2Ks that then phosphorylate MAPKs JNK and p38α, and is a sensor for many different ribotoxic stressors ^1^, the possibility that it might be activated by 2-5A-induced RNase L activity was investigated. Remarkably, phosphorylation of JNK and p38α in response to 2-5A and RNase L was abrogated in ZAK KO BMM (Fig. 1F). There was, however, no effect of ZAKα on the ribonuclease activity of RNase L (Fig. 1F, lower panel). Furthermore, a small molecule ZAK inhibitor (compound 3h, ZAKi) which blocks the kinase activity of ZAKα ^28^, greatly reduced phosphorylation of both JNK and p38α in response to 2-5A, without affecting the ribonuclease activity of RNase L (Fig. 1G). These data show that ZAKα is required for RNase L-induced JNK and p38α phosphorylation in a kinase-dependent manner during the ribotoxic stress response.

### 2-5A activation of RNase L modulates the macrophage transcriptome by degrading and inducing different transcripts

To probe 2-5A-induced global effects on the transcriptome, WT BMMs and RNase L KO BMMs were either mock transfected or transfected with 2-5A or A3 followed by bulk RNA-sequencing (Fig. S2A). Transcriptomes of the 2-5A-transfected WT BMM were significantly different from all other treatment groups, indicating that RNase L activity dramatically altered transcript expression (Fig. 2A). The differentially expressed genes (DEGs) for every condition were compared and cross-analyzed (Figs. 2B, S2C-F). There were 4843 DEGs in 2-5A-stimulated WT BMMs, comprising about 20% of the total transcripts sequenced (Fig. 2B and Table S2). In contrast, there were no DEGs in A3-transfected WT BMMs nor in 2-5A- or A3-transfected RNase L KO BMMs. Therefore, transcriptomic alterations in 2-5A-stimulated WT BMMs were exclusively due to RNase L activity. Among these DEGs were 3,334 genes with reduced expression (presumably due to RNA degradation) and 1,509 genes with increased expression (Fig. 2C). Given that RNase L is a general ribonuclease with only modest specificity, the observation that a minority of transcripts were decreased in abundance suggested that some transcripts might be preferentially degraded by RNase L ^19^. These results are in contrast with studies showing 70% to 90% mRNA degradation in dsRNA(pIC)-transfected cells ^21,23^. The differences in the extent of RNA degradation are likely due to differences in the level and duration of RNase L activation between the two types of protocols, i.e. 2-5A-transfection or pIC-transfection, and to differences in expression levels of RNase L in the different cell types.

**Figure 2.**
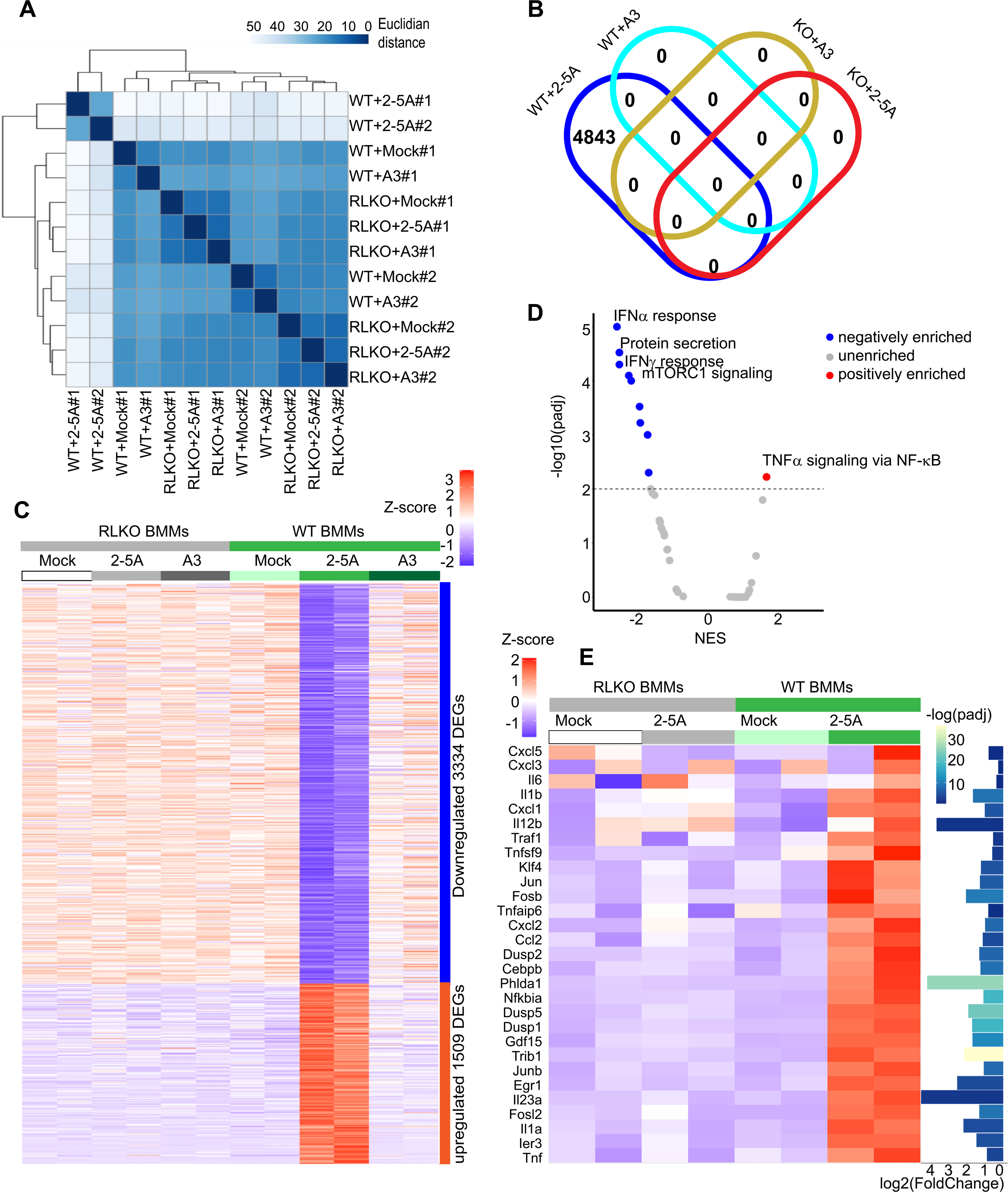
RNase L activation by 2-5A modulates the transcriptome of BMM, decreasing or increasing levels of different mRNAs. (A) Euclidean distance of gene expression between treatment groups as a measure of sequence divergence^56^. The matrix of distances between each pair of samples is represented by a dendrogram, while the scale of sample-to-sample distances is represented in different shades of blue. (B) Shared DEGs in mock-, 2-5A- or A3-transfected WT or RNase L KO BMMs. The Venn diagram depicts transcript changes shared or unique between each comparison. (C) Heatmaps of mock-, 2-5A, and A3-transfected WT and RL KO BMM. The color bar (upper right) indicates the Z-score. (D) Volcano plot of 2-5A down- and up-regulated gene sets from WT BMMs transcriptomes. (E) Heatmap of “TNF-α signaling via NF-κB” Hallmark gene set from mock- and 2-5A-transfected WT and RL KO BMM. Inserts: Z-score (upper left) and adjusted *p* values (padj) (right insert). RL KO, RNase L KO.

To better understand the functional impact of 2-5A-induced RNase L activity on the transcriptome, gene set enrichment analysis (GSEA) was performed based on the DEG ranking from 2-5A-transfected WT BMM RNAseq. Among Hallmark Gene Sets, which classify genes according to their participation in biological states or processes, there were predominantly decreases (negative enrichment) of transcripts related to gene sets corresponding to IFNα and IFNγ responses, protein secretion and mTORC1 signaling (Fig. 2D). In contrast, there was a relative increase (positive enrichment) of transcripts in the “TNFα signaling via NF-κB” gene set, including many genes encoding proinflammatory proteins (IL6,TNF) and transcription factors (Jun,Fosb) (Fig. 2D&E). In addition, GSEA Gene Ontology (GO) analysis based on functions of genes and gene products indicated that RNase L predominantly reduced levels of certain transcript groups, including for antigen processing and presentation, small ribosomal subunit, IFNα production, regulation of defense to virus, and the innate immune response (Fig. S3A). Effects of 2-5A/RNase L on transcripts for small ribosomal subunit proteins could possibly contribute to ribosomal stress. On the other hand, 2-5A stimulation positively enriched acute phase response transcripts, including proinflammatory genes (Fig. S3A).

Downregulated transcripts from 2-5A-transfected WT BMM are represented in heatmaps for IFN stimulated genes (ISGs) and antiviral genes (Fig. S3B), antigen processing and presentation (Fig. S3C), ribosomal protein genes (Fig. S4A) and housekeeping genes (Fig. S4B). In accord with GSEA analysis, the proinflammatory genes downstream of TNFα, especially those related to stress responses, were induced in WT BMMs upon 2-5A stimulation (Fig. 2E). In addition, we confirmed 2-5A induction of the growth differentiation factor 15 gene (Gdf15) and added it to the heatmap(Fig. 2E)^29^. These data indicate that 2-5A-stimulated RNase L activity led to bidirectional changes in levels of different transcripts, increases in proinflammatory and stress-responsive transcripts but decreases in many other transcripts, including those involved in IFN responses, antigen processing and presentation, and small ribosome subunit proteins.

### 2-5A/RNase L induces a proinflammatory transcriptome signature in BMM through *de novo* mRNA synthesis

To validate the GSEA, levels of representative transcripts from different categories of genes were monitored by qRT-PCR. Transcript that decreased in relative amounts after 2-5A treatment of WT BMM included those in IFN responses (Ifit1bl1, Stat1, Irf7, Gbp2, Rsad2), inflammation (Cxcl9) and antigen processing and presentation (H2-Aa1, H2-Eb1, Cd86). Transcripts that increased in abundance in 2-5A-transfected WT BMM included TNFα-regulated genes and stress-response genes (Cxcl2, Gdf15, IL-1β, IL-23α) (Fig. 3A). These changes were not observed in 2-5A transfected RNase L KO BMMs. Results were normalized to GAPDH mRNA levels (Fig. 3A). Because RNase L activation decreased mRNA levels for housekeeping genes, including GAPDH mRNA, transcript levels were separately normalized by input mRNA levels (Figs. 3B and S4B). Nevertheless, RNA levels resulted in similar expression patterns when normalizing to levels of input mRNA (Fig. 3B). Those results suggested positive enrichment was due to increases in levels of transcripts related to proinflammatory responses.

**Figure 3.**
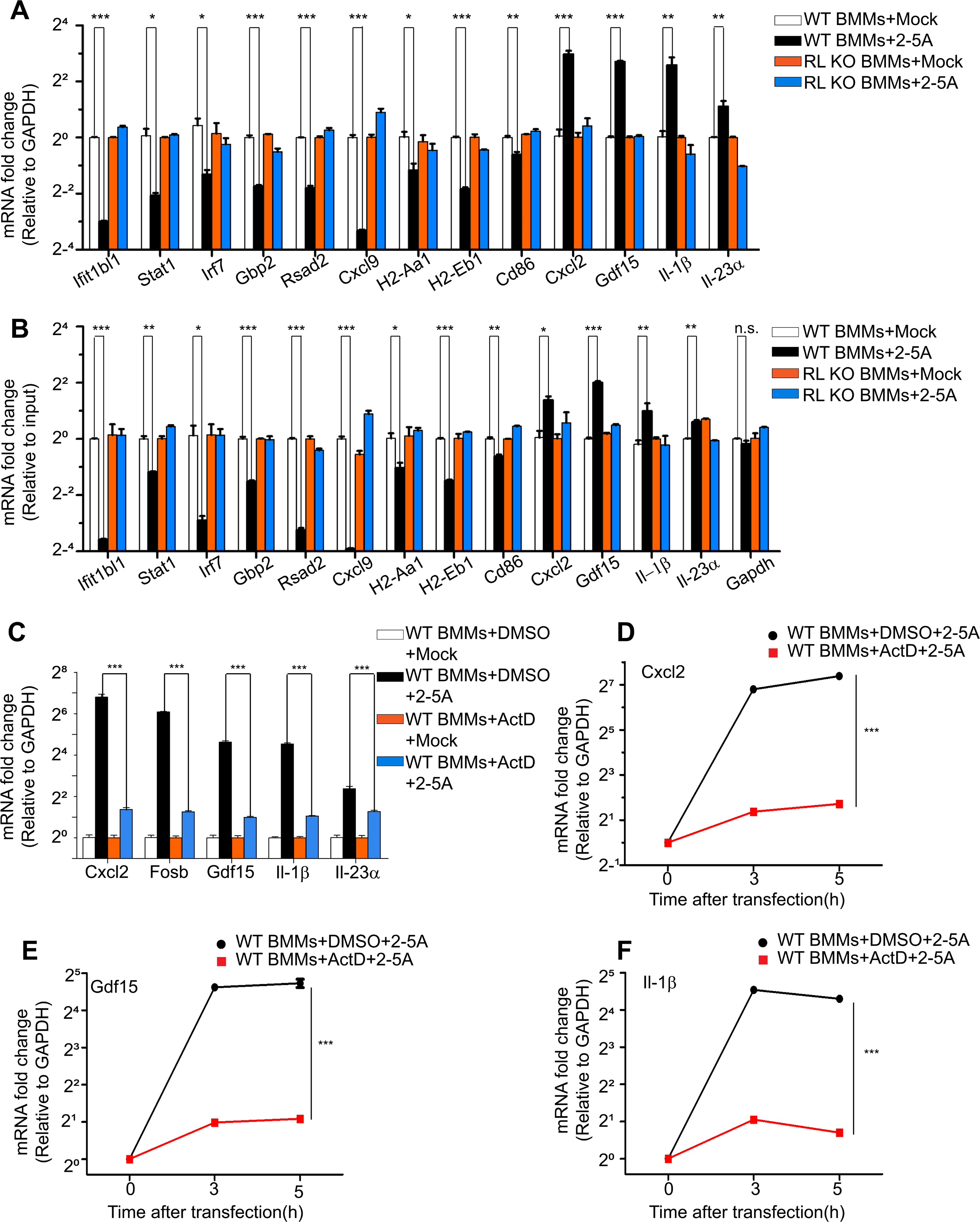
Regulation of transcript levels in mock- or 2-5A-transfected WT and RL KO BMM. (A, B) Levels of select up- and down-regulated transcripts in mock- and 2-5A-transfected (20 μM for 3 h) WT and RL KO BMM normalized to (A) GAPDH mRNA levels or to (B) input RNA levels as determined by qRT-PCR. (C-F) Actinomycin D inhibited 2-5A-induction (20 μM for 3 h) of select transcripts in WT BMM as determined by qRT-PCR. Actinomycin D (5 μg per ml) or DMSO, was added to cells 30 min prior to transfections. Relative transcript levels were monitored by qRT-PCR. Experiments performed with three replicates were repeated at least twice. Significance was determined by unpaired Student t-test. ***, *p*<0.001; **, p<0.01; *, *p*<0.05. ActD, actinomycin D; RL KO, RNase L KO.

To determine if the 2-5A/RNase L-induced transcripts reflected transcriptional induction or resistance to RNase L ribonuclease activity, WT BMMs were treated with actinomycin D to block transcription during 2-5A transfection. Indeed, actinomycin D greatly inhibited the increases in transcript levels suggesting transcriptional induction in response to 2-5A (Fig. 3C-F). Current findings are consistent with the observation of transcriptional-induction of the Gdf15 gene in response to 2-5A activation of RNase L ^29^ (Figs. 2E and 3E).

DsRNA has many effects on cells besides OAS activation (notably on MAVS/IRF3 signaling and PKR activation). To compare and contrast effects of dsRNA and 2-5A on gene regulation, selected transcripts were monitored by qRT-PCR in WT BMM transfected with poly(I):poly(C) (pIC) or 2-5A. Results showed that levels of some pIC-induced transcripts were decreased following 2-5A transfection, likely due to RNase L-mediated mRNA degradation (Fig. 4A-H). On the other hand, some proinflammatory genes induced by 2-5A (Cxcl2, IL-1β and IL-23α), were also induced upon pIC stimulation in WT BMM (Fig. 4I-K). These results emphasize why it was necessary to use 2-5A to identify transcripts that are specifically regulated by RNase L.

**Figure 4.**
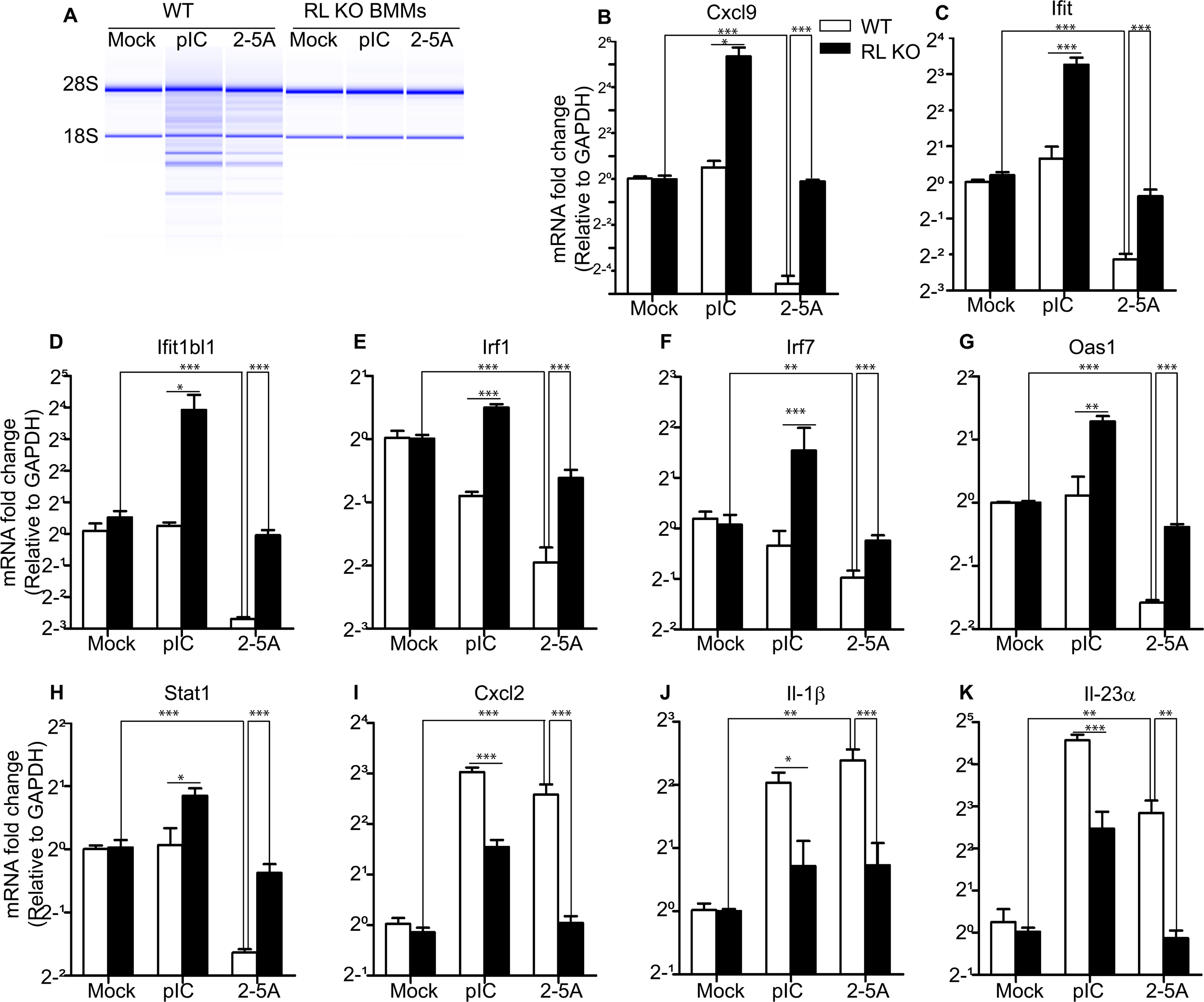
Comparison of representative dsRNA- and 2-5A-induced or -repressed transcripts. (A) Cleavage of rRNA in WT BMM, but not in RNase L KO BMM, in response to poly(I):poly(C) (pIC) (1 μg per ml, 3 h) or 2-5A (20 μM, 3 h) as determined by analysis on an RNA chip (Agilent). (B-K). Relative levels of different transcripts (as indicated) with transfection of pIC (1 μg per ml, 3 h) or 2-5A (20 μM, 3 h) as determined by qRT-PCR. ***, *p*<0.001; **, p<0.01.

### The ZAKα-initiated MAP kinase cascade induces inflammatory gene transcripts in response to 2-5A

To investigate how proinflammatory genes were induced by 2-5A, three different pathways were considered: (1) NF-κB mediated transcription suggested by Gene Ontology analysis (Fig. S3A); (2) the MAVS-IRF3 pathway suggested by our previous study ^30^, and (3) the JNK/p38α pathway because of its involvement in ribotoxic stress responses and its activation during TNFα signaling ^6^. However, NF-κB was not activated by 2-5A stimulation, and inhibition of the NF-κB upstream kinase, IKK-β, did not affect levels of 2-5A-induced transcripts in WT BMM (Fig. S5A,B). In addition, 2-5A-induction of transcripts for Cxcl2, IL-1β and IL-6 was not prevented by deletion of MAVS in A549 cells (Fig. S5C). In contrast, MAP kinases JNK and p38α and their downstream transcription factor AP-1 component c-Jun were phosphorylated upon 2-5A stimulation of WT BMM (Figs.1E and 5A) ^31^. The inhibitor of p38α (p38αi) prevented 2-5A induction of Cxcl2, Fosb, Gdf15, and IL-23α but not IL-1β (Fig. 5B). The inhibitor of JNK (JNKi) reduced 2-5A induction of all transcripts except IL-23α (Fig. 5C), whereas inhibitor of AP-1 (AP-1i), a transcription factor downstream of JNK/p38α, decreased induction of all transcripts examined (Fig. 5D). These results suggest that both p38α and JNK were involved in 2-5A induction of transcription for at least some of the proinflammatory genes.

**Figure 5.**
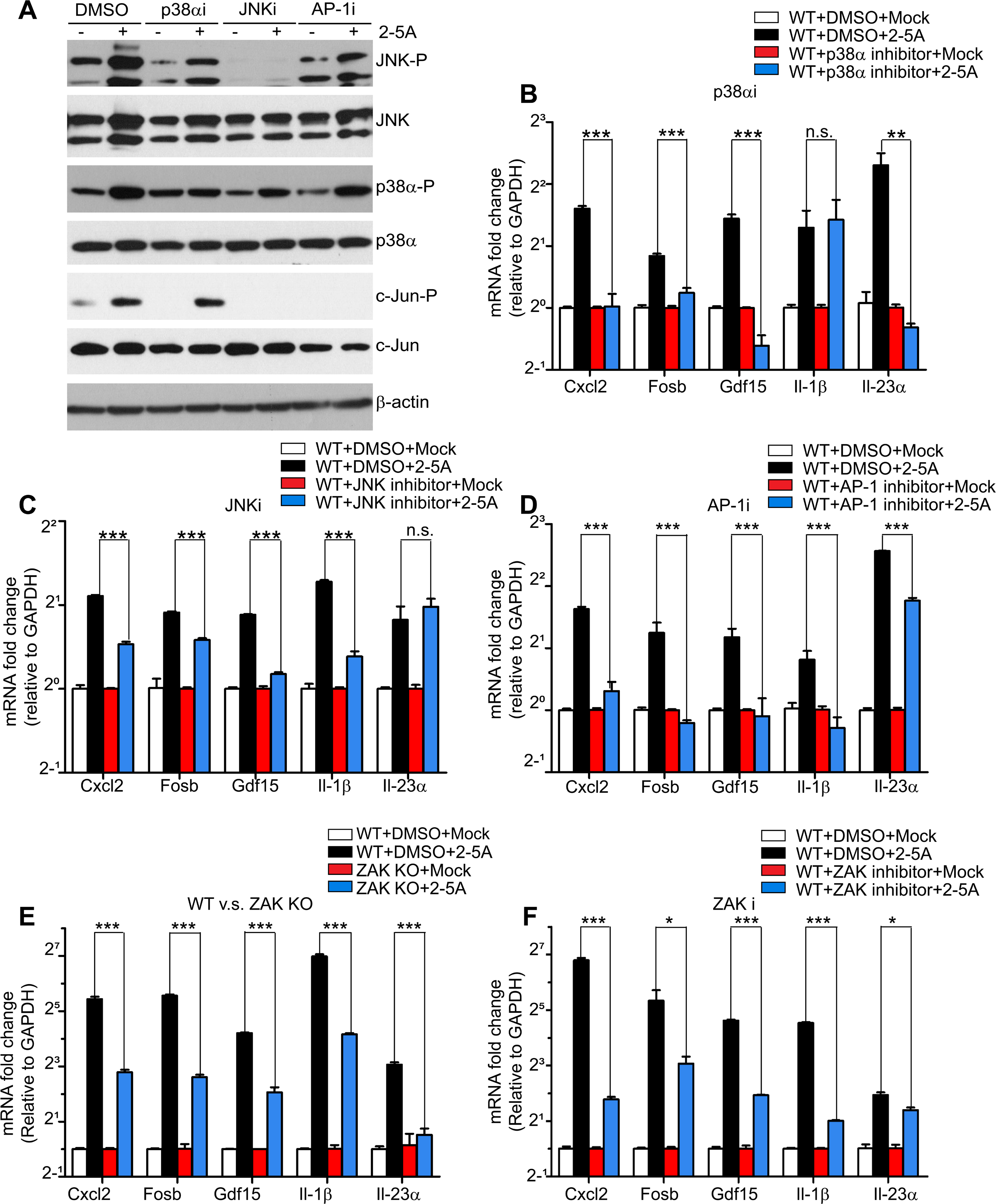
Effects of p38α, JNK, AP-1 and ZAK inhibitors or ZAK knockout on 2-5A induction of transcripts in WT BMM. (A) Inhibition of p38α, JNK, and c-Jun phosphorylation by small molecule inhibitors in 2-5A-transfected WT BMM as determined by Western blotting. Inhibitors, or DMSO, were added to cells for 2 h prior to transfections with 2-5A (20 μM for an additional 3 h). (B-D) Effects of inhibitors of (B) p38α, (C) JNK, and (D) AP-1 on transcript levels after 2-5A transfection in WT BMM. (E) 2-5A induction (20 μM, 3 h) of transcript levels in WT and ZAK KO BMM. (F) Effects of ZAK inhibitor (compound 3h) on induction of transcript levels in WT BMM. Significance was determined by unpaired Student t-test. ***, *p*<0.001; **, *p*<0.01; *, *p*<0.05.

Because ZAKα is an upstream kinase for JNK and p38α phosphorylation, WT BMM and ZAK KO BMM were compared for effects of 2-5A on gene induction. Levels of all five transcripts examined were significantly reduced in ZAK KO BMM compared to the WT BMM (Fig. 5E). Similarly, a ZAK inhibitor (compound 3h, ZAKi) reduced 2-5A induction of all five transcripts in WT BMM (Fig. 5F). These results suggest that ZAKα is required for 2-5A induction of these proinflammatory genes in a kinase activity dependent manner.

### ZAKα is required for 2-5A/RNase L regulation of transcript levels and for JNK and p38α phosphorylation in human monocytes

To extend these findings to a human immune cell type, ZAK was knocked out in the human monocytic cell line THP-1. The human homologs of some representative genes from the BMM studies were examined in the THP-1 cells with similar observed changes upon 2-5A transfection when normalized to SON DNA and RNA binding protein (SON) mRNA, which is resistant to RNase L cleavage (Fig. 6A,B) ^23^. Transcripts for Cxcl2, Fosb, GDF15, IL-1β, and IL-23α were induced by 2-5A transfection of WT THP-1 cells, but not in RNase L KO THP-1 cells (Fig. 6A).

**Figure 6.**
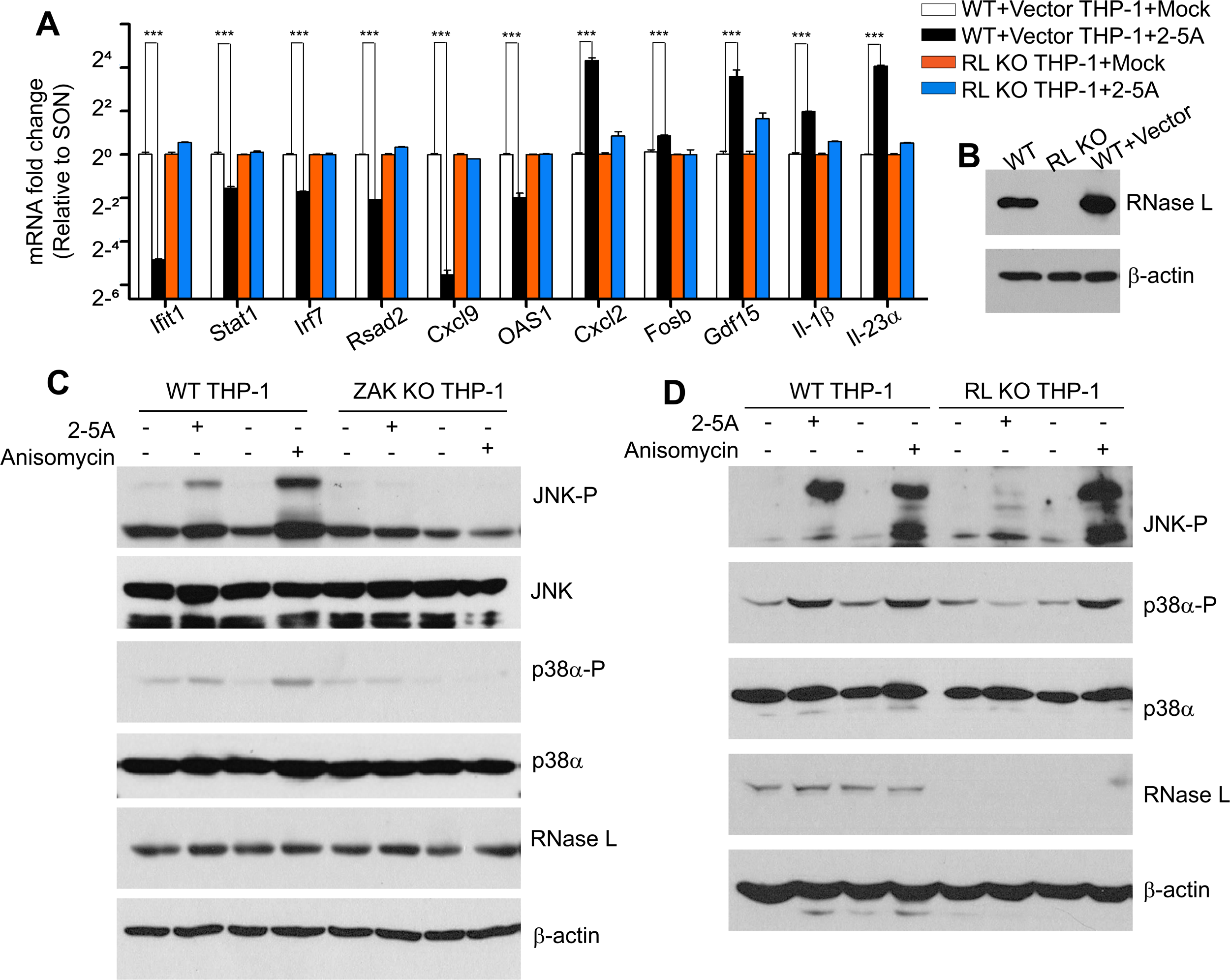
ZAKα is required for the ribotoxic stress response in human THP-1 monocytic cells. (A) Transcript levels in mock- and 2-5A-transfected (20 μM for 3h) WT and RL KO THP-1 cells normalized to SON DNA and RNA binding protein (SON) mRNA levels. Data are from three separate experiments, each performed three times. Significance was determined by unpaired Student t-test. ***, *p*<0.001. (B) Western blot of WT, RL KO, and WT vector control THP-1 cells probed with antibody against human RNase L or β-actin. (C,D) Levels of total and phosphorylated JNK and p38α, and of RNase L and β-actin in (C) WT and ZAK KO THP-1 cells, or (D) WT and RL KO THP-1 cells that were mock-transfected, or transfected with 2-5A (20 μM, 2 h) or treated with anisomycin (5 μg per ml, 1 h) as determined in Western blots. RL KO, RNase L KO.

Furthermore, both 2-5A- and anisomycin-induced JNK/p38α phosphorylation was impaired in THP1 cells lacking ZAK (Fig. 6C). In contrast, while RNase L knockout prevented 2-5A-induced phosphorylation of JNK and p38α, the absence of RNase L had no effect on anisomycin induced activation of JNK and p38α. These results show that 2-5A/RNase L specifically regulates ZAKα signaling during the ribotoxic stress response in both human monocytes and mouse bone-marrow derived macrophages.

### Apoptosis of cancer cells in response to 2-5A/RNase L requires ZAKα

Because evasion of apoptosis is a hallmark of cancer ^32^, the possible role of ZAKα in apoptosis triggered by 2-5A/RNase L was investigated in human lung adenocarcinoma A549 cells. To determine if ZAKα was phosphorylated in response to 2-5A, gel mobility shift assays were done with Phos-Tag gels. The slower migrating phosphorylated ZAKα is thereby separated from unphosphorylated ZAKα ^4^. Both 2-5A or anisomycin caused ZAKα auto-phosphorylation as determined by the mobility shift. However, anisomycin was more effective than 2-5A in causing phosphorylation of ZAKα, p38α and JNK (Fig. 7A). In addition, either of two ZAK inhibitors (compounds 3h & 6p) ^28,33^ prevented phosphorylation of ZAKα and its downstream MAPKs, JNK and p38α, in response to either 2-5A or anisomycin. These results show that ZAKα kinase activity is necessary for 2-5A-induced JNK and p38α activation.

**Figure 7.**
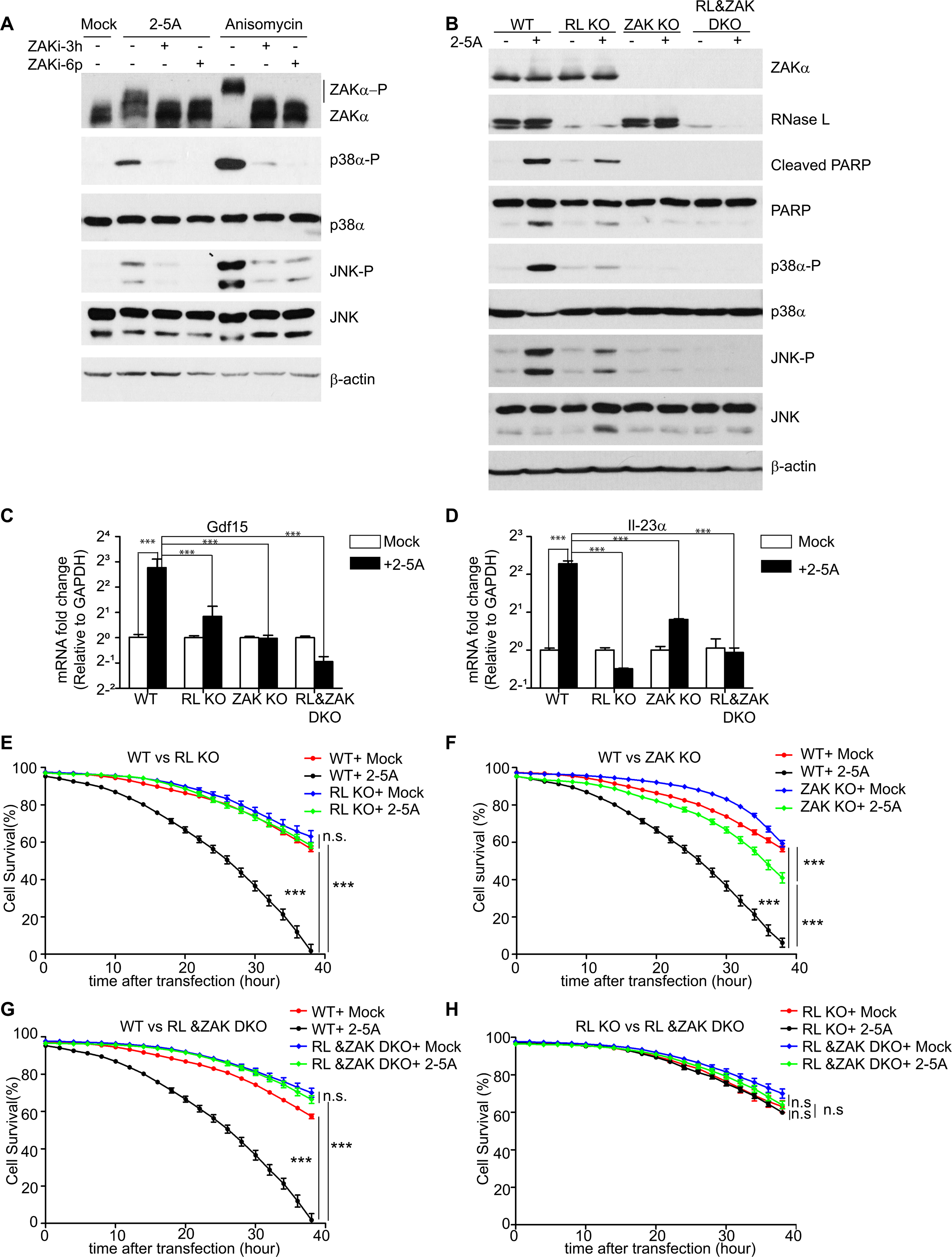
Involvement of ZAKα in JNK/p38α phosphorylation, gene regulation and apoptosis triggered by 2-5A activation of RNase L in A549 cells. (A) ZAKα phosphorylation in response to 2-5A transfection (20 μM, 2 h) or anisomycin (5 μg per ml, 1 h) in the presence or absence of ZAKα inhibitor compounds 3 h (10 μM) or 6 h (10 μM) of WT A549 cells as determined by gel mobility shift assay. Western blots were also probed with antibodies against total and phosphorylated p38α and JNK, and β-actin. (B) Western blots for total and cleaved PARP and total and phosphorylated p38α and JNK in mock-transfected and 2-5A-transfected (20 μM, 16 h) WT, RL KO, ZAK KO and RL&ZAK DKO A549 cells. (C,D). Levels of (C) GDF15 and (D) IL-23α transcripts relative to GAPDH transcript levels in WT, RL KO, ZAK KO and RL&ZAK DKO A549 cells after mock- or 2-5A-transfection (20 μM, 3 h) as determined by qRT-PCR. [Results were reproduced in two separated experiments, each in triplicate]. Significance was determined by unpaired Student t-tests. (E-H) Cell survival curves for WT, RL KO, ZAK KO, RL&ZAK DKO A549 cells (as indicated) that were mock- or 2-5A-transfected (20 μM 2-5A). For comparisons between different treatment groups, the same reference data sets for WT, RL KO, and RL&ZAK DKO are included in different panels. Cell viability was determined by real-time imaging of cells simultaneous stained for total cells and dead cells. Significance was determined by two-way ANOVA. ***, *p*<0.001; ns, not significant.

To investigate the involvement of ZAKα and RNase L in apoptosis of A549 cells, these proteins were deleted from A549 cells singly (KO) and in combination (i.e., double knockout or DKO). RNase L knockout inhibited both PARP cleavage and JNK/p38α phosphorylation upon 2-5A transfection. Similarly, 2-5A-induced JNK/p38α phosphorylation and PARP cleavage was inhibited in ZAK KO and in ZAKα/RNase L DKO cells (Fig. 7B). Also, 2-5A induction of Gdf15, IL-23α, Cxcl2, Fosb, and IL-1β transcripts were inhibited in RNase KO, ZAK KO and in ZAK/RNase L DKO cells (Figs. 7C,D and S6A-C).

Cell death in response to 2-5A was monitored by cell staining and real-time imaging of cells. 2-5A transfection of WT A459 cells resulted in a dramatic reduction in cell survival (Figs. 7E-G and S6D). In contrast, in both RNase L KO and RNase L/ZAK DKO there was no increase in cell death after 2-5A transfection (Fig. 7E,G,H). Deletion of ZAKα alone greatly inhibited cell death (by about 50%) in response to 2-5A, consistent with involvement of ZAKα in RNase L-mediated cell death (Fig. 7F). However, there was a smaller increase in cell death (about 20%) after 24 h in response to 2-5A in ZAK KO cells, possibly indicating an alternative, non-apoptotic pathway of 2-5A/RNase L induced cell death as there was no detectable PARP cleavage (Figs. 7B and F). These data indicate that ZAKα is the predominant mediator of cell death signaling in response to activation of RNase L in a human cancer cell line.

## DISCUSSION

### Impact of RNase L on innate immune signaling

Here we show that RNA degradation by RNase L triggers a kinase cascade that leads to innate immune signaling and apoptosis. These events are set in motion when dsRNA binds and activates OAS enzymes that produce the unusual allosteric effector of RNase L, 2-5A. Innate immune signaling is often initiated by the presence of dsRNA in the cytoplasm. Many RNA and DNA viruses produce non-self dsRNA, as intermediates or by-products of their infection cycles. However, even in the absence of viral infections cells can sometimes produce self dsRNA capable of trigger innate immune responses. For instance, self dsRNA is produced by transcription of repetitive DNA elements in the genome in response to promoter hypomethylation ^10,34^. In addition, genetic deficiency of the dsRNA editing enzyme, ADAR1, leads to accumulation of host dsRNA ^35^. In both instances there is activation of the 2-5A/RNase L pathway. DsRNA principally activates three types of innate signaling pathways. (1) Upon sensing dsRNA, RIG-I like receptors (RLRs) signal through MAVS to IRF3 and NF-κB to induce transcription of genes for IFNs and other pro-inflammatory cytokines ^36^. (2) The IFN-inducible and dsRNA-dependent protein kinase, PKR, phosphorylates eIF2α causing inhibition of most protein synthesis ^37^. (3) DsRNA stimulates OASs to produce a series of unusual 5’-triphosphorylated 2’,5’-linked short oligoadenylates (p_3_(A2’p5’)_n≥2_A or 2-5A) that activate RNase L ^9,12,13,17^. By cleaving ssRNA, RNase L has profound effects on cells. For instance, RNase L activation beyond a threshold level cause the infected cells to spiral into apoptosis, preventing further virus replication ^22,38,39^. RNase L-deficient mice have a compromised ability to survive various types of viral infections, including West Nile virus, coxsackievirus B4, and encephalomyocarditis virus (EMCV) ^39–41^. OAS-RNase L inhibits many types of RNA viruses, including SARS-CoV-2 (but not Zika virus where RNase L actually increases viral replication) and at least one type of DNA virus, the poxvirus vaccinia virus ^11,42–44^. In this context, our study reveals a pathway by which RNase L upregulates transcript levels for proinflammatory genes that typically restrict viral infections.

### Degradation and induction of different RNAs by 2-5A activation of RNase L

In recent years, insight into complex biological processes has been obtained through quantitative and comprehensive examination of the RNA pool in cells (RNAseq) by next-generation sequencing. Characterizing the host innate immune response to infection by pathogens is one of many areas in which RNAseq is informative ^45^. Virus infection has profound effects on host cell gene expression owing in part to induction of IFNs and their downstream effects. In particular, the OAS-RNase L pathway is responsible for the rapid degradation of viral and host ssRNA ^17,21,23,46^.

Dissecting effects of RNase L on the transcriptome of virus-infected cells is complicated in part because viruses typically replicate to higher levels in RNase L KO cells than WT cells ^17,39^. Therefore, virus-induced alterations in the transcriptome may be amplified in RNase L KO cells, thus obscuring direct effects of RNase L on RNA levels. For instance, viruses induce type I and III IFNs which lead to expression of hundreds of IFN stimulated gene (ISG) transcripts (Der et al., 1998). An alternative approach of transfecting cells with dsRNA, typically the synthetic dsRNA, poly(I):poly(C) (pIC) as a virus surrogate, has its own fingerprint on the transcriptome. Among the effects of dsRNA are induction of IFNs and other cytokine mRNAs, or their degradation, as well as OAS-RNase L and PKR activation^47^. Therefore, in the present study we directly and specifically activated RNase L in cells with its natural ligand 2-5A.

There were the expected decreases in levels of a wide range of different gene transcripts by RNase L mediated RNA cleavage, but there was also induction of many transcripts for proinflammatory cytokines, chemokines, and transcription factors. These present results confirm our prior report of 2-5A-induced upregulation of many proinflammatory mRNA species, including [IL8 (CXCL8), CXCL1, IL17R] ^29^. The 2-5A/RNase L induced genes are consistent with a ribotoxic stress response through JNK and p38α ^22,26^. The results provide an intriguing level of complexity to the regulation of the transcriptome by RNase L.

### RNase L activation and the ribotoxic stress response

Ribotoxic stress responses are cellular mechanisms for coping with agents or treatments that cause ribosome stalling and collision ^1,3,4^. Ribotoxic stressors often damage RNA either directly by ribonucleases (α-sarcin, ricin) or by agents (uv, doxorubicin) that damage RNA in other ways ^48^. Here we report that activation of RNase L by its specific allosteric effector, 2-5A, leads to a ribotoxic stress response requiring ZAKα for JNK and p38α phosphorylation resulting in inflammatory signaling and apoptosis (Fig. 8). The same mechanism was observed in mouse BMM, human THP-1 and A549 cells. Interestingly, induction of proinflammatory transcripts by 2-5A/RNase L occurs simultaneously with widespread massive degradation of ssRNA. Our transcription factor inhibitor studies suggest the induced genes were likely the result of JNK and p38α phosphorylation leading to activation of AP-1 transcription factor.

**Figure 8.**
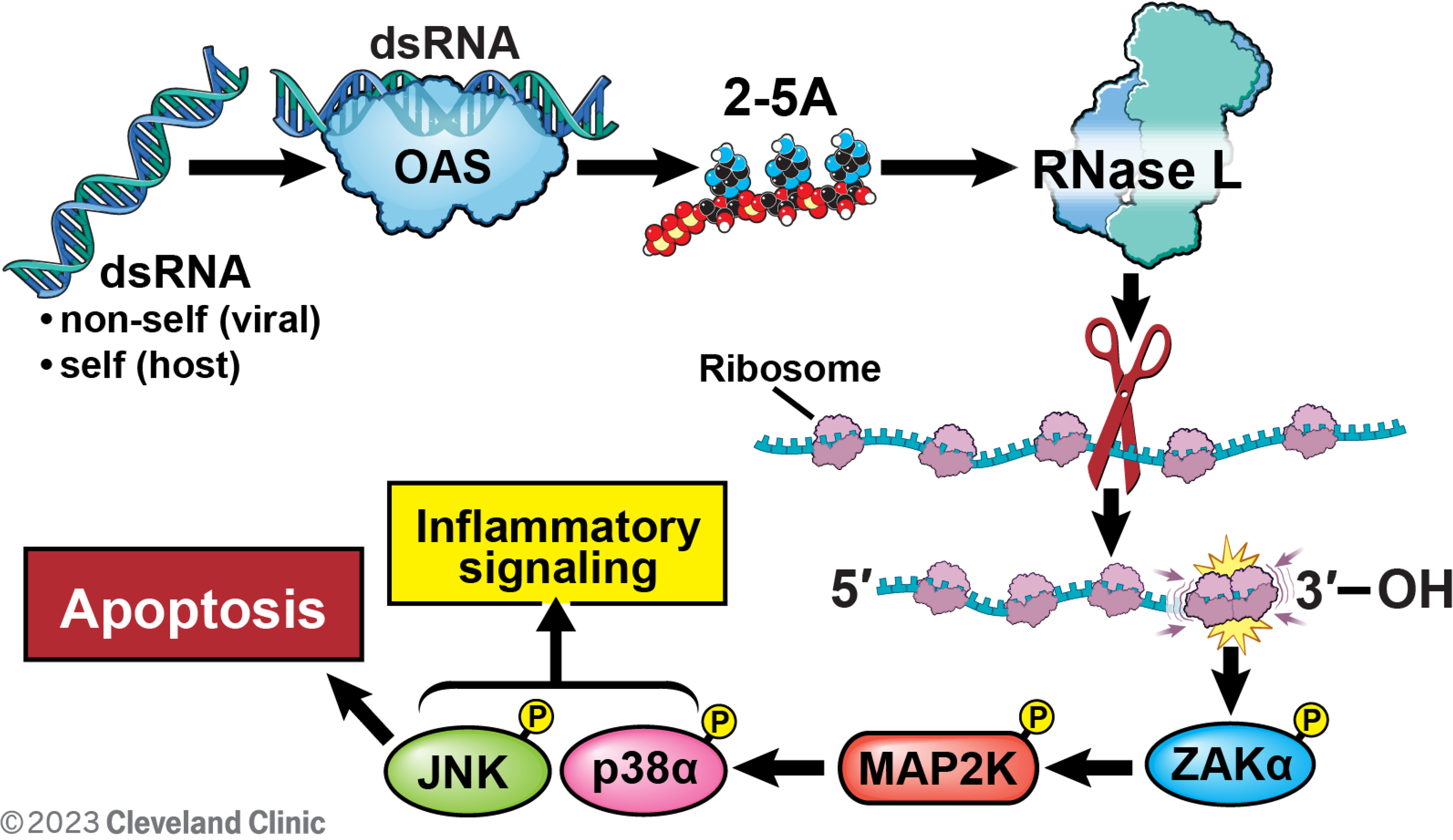
The ribotoxic stress response mediated by the OAS-RNase L pathway through ZAKα signaling. OASs 1-3 produce 2-5A in response to virus dsRNA or host dsRNA. 2-5A activates RNase L which cleaves mRNA in polysomes. Ribosomes collisions then occurs at the 3’ ends of cleaved RNAs lacking a stop codon, triggering ZAKα activation. Subsequently, ZAKα phosphorylates MAP2Ks that phosphorylate JNK and p38α leading to inflammatory signaling and apoptosis.

Although RNase L cleaves rRNAs and some tRNAs, it is cleavage of mRNAs that is believed to primarily impair protein synthesis ^20,21,23–25^. There is a net accumulation of some proinflammatory mRNAs in response to 2-5A activation of RNase L suggesting that for these transcripts mRNA synthesis rates exceeded degradation rates. Because rRNA degradation by RNase L does not appear to inhibit translation there may not be an impediment to the translation of the induced mRNAs ^20,23^. Accordingly, we previously showed that 2-5A/RNase L induced transcription of the *GDF15* gene resulted in production of the GDF15 protein ^29^.

Our results support a role for the OAS-RNase L in regulating inflammation and apoptosis in response to viral (non-self) dsRNA, or host (self) dsRNA (Fig. 8) ^10,35,46,49^. 2-5A produced from dsRNA-stimulated OASs activates RNase L ^9,17^. While precisely how RNase L activity leads to ZAKα activation is unknown, prior studies suggest a mechanism. Activation of ribosome-associated ZAKα occurs in response to ribosome stalling and collisions ^4,50^. In addition, RNase L cleaves mRNAs internally in actively translating polysomes resulting in translation of alternative ORFs present in the RNA cleavage products derived internally to the coding sequence and in the 5’- and 3’-UTRs^51^. When translating RNAs lack a stop codon, ribosomes do not dissociate, likely resulting in ribosome collisions ^52^. Therefore, a potential pathway to ZAKα activation is ribosome collisions on RNase L-mediated RNA cleavage products lacking a stop codon (Fig. 8). It remains to be determined, however, whether RNase L activation is associated with local and/or widespread ribosome collision. Auto-phosphorylation of the ribosome-binding C-terminus of ZAKα causes the activated kinase to dissociate from ribosomes and then to activate MAP2Ks ^3,6^. MAP2Ks then phosphorylate JNK and p38α, resulting in inflammatory signaling and apoptosis. Therefore, the ribotoxic stress response provides a mechanistic framework for understanding of how RNase L activation by 2-5A leads to induction of proinflammatory genes and apoptosis. Future studies on OAS-RNase L activation of ZAKα might therefore inform drug development efforts aimed at controlling cancer and virus-mediated inflammation.

## MATERIALS AND METHODS

### Reagents

Trimer 2-5A (ppp5’A2’p5’A2’p5’A) was enzymatically synthesized from ATP with histidine-tagged porcine OAS1 and was HPLC-purified as we described previously^53^. A3 (A2’p5’A2’p5’A) was prepared by dephosphorylation of 2-5A as described ^53^. Poly(I):poly(C) (pIC) was purchased from Millipore Sigma(#528906). The small molecule inhibitors used in this study were JNK inhibitor (SP600125, Santa Cruz, #sc-200635); p38a inhibitor (SB203580, Millipore Sigma, #559389); AP-1 inhibitor (SP100030, Sigma-Aldrich, #5315350001); IKKβ inhibitor (BAY11-7082, Santa Cruz, sc-200615B); and ZAK inhibitors (3h and 6p) ^28,33^. Anisomycin was from Sigma-Aldrich, #A9789. The antibodies from Cell Signaling Technology were phosphor-p38a antibody (Thr180/Tyr182, #4511); p38a total antibody (#9212s); phospho-JNK antibody (#9255 and #4668), SAPK/JNK antibody (#9252 and #9258), phospho-p44/42 ERK1/2 antibody (Thr202/Tyr204) (#9101), p44/42 MAPK ERK1/2 antibody (#4695), PARP total antibody(#9542), and cleaved PARP (Asp214) antibody (#5625). Anti-mouse RNase L (R-188, rabbit polyclonal) antibody was produced in-house (B.K. Jha and R.H. Silverman), and anti-human RNase L (mouse monoclonal antibody) was previously described ^14^. Anti-mouse ZAKa antibody (proteintech, #14945-1-AP) and anti-human ZAKa antibody were from Thermo Fisher Scientific (A301-993A). Anti-a-tubulin was from Sigma-Aldrich (#T9026).

### Mice

*Rnasel*^-/-^ (KO) mice were previously generated and back-crossed on a C57BL6 background ^39,54^. Wild type (WT) C57BL6 mice were purchased from The Jackson Laboratory. All mouse studies at Cleveland Clinic were performed in compliance with a protocol approved by the Institutional Animal Care and Use Committee of the Cleveland Clinic Lerner Research Institute. ZAK KO mice in C57BL/6 background ^50^ were housed in the animal facility of the Department of Experimental Medicine at the University of Copenhagen under the oversight of the Institutional Animal Care and Use Committee. All mouse experiments were conducted in compliance with Danish regulations and approved by the Danish Animal Experiments Inspectorate. Mice were maintained on a 12-h light:dark cycle and had unrestricted access to commercial rodent chow and water prior to the experiment.

### Cell culture

THP-1 cells obtained from ATCC (TIB-202) were cultured in RPMI1640 medium (Media Preparation Core, Lerner Research Institute) supplemented with 10% FBS (Gibco Ref: 10437-028 from Thermo Fisher Scientific) and 50 μM of β-mercaptoethanol (Sigma-Aldrich). A549 cells obtained from ATCC (CCL-185) were cultured in RPMI1640 medium supplemented with 10% FBS. *Rnasel* KO of cell lines was done with a CRISPR-cas9 lentivirus generated by transfecting the LentiCRISPER-sgRL-6 [a gift from Dr. Susan Weiss, University of Pennsylvania ^11^] or control LentiCRISPR Vector 2, with psPAX2 and pVSV-g plasmids at 2:1:1 ratio into 293T cells cultured in DMEM medium (Media Preparation Core, Lerner Research Institute) supplemented with 10% FBS. ZAK knockout THP-1 and A549 cells were done with a CRISPR-cas9 lentivirus by transfecting the LentiCRISPER-sg (the gRNA sequence was from ^3^) or control LentiCRISPR Vector 2, with psPAX2 and pVSV-g plasmids at 2:1:1 ratio into 293T cells. The virus supernatant was harvested 72 h after transfection and passed through a 0.22 μm filter. Infections were done on THP-1 or A549 with 1 mL virus supernatant supplemented with 8 μg/mL polybrene. Cells were centrifuged in 6-well plates at 600xg for 2 h and cultured overnight. Subsequently, cells were collected and centrifuged at 300xg for 5 min. Cells were resuspended into RPMI1640 medium (supplemented with 10% FBS) containing puromycin (2 μg/mL) and pipetted into 96-well plates for selection of single colonies. After 4 weeks of selection, surviving colonies were validated by probing Westerns blots for absence of RNase L with monoclonal antibody against human RNase L ^14^. MAVS KO and its control A549 cells (all cultured in DMEM with 10% FBS) were described previously and were gifts from Susan R. Weiss (University of Pennsylvania) ^35,55^.

### Bone marrow derived macrophages

Bone marrow cells were isolated from hind limbs of age and gender matched 8- to 12-week-old WT or *Rnasel*^-/-^ mice. After removing the red blood cells (Gibco, A10492-01), remaining cells were seeded at 4×10^6^ cells per well in 6-well plate in 2 mL of DMEM medium supplemented with 10% FBS, 50 μM β-mercaptoethanol and 20 ng/mL recombinant murine M-CSF (315-02, Peprotech) with medium changed every 2 days. After 7 days of culture, cells were ready for transfection.

### Cell transfections or stimulation

WT or *Rnasel* KO macrophages were pretreated for 1 h with JNK inhibitor (30 μM), p38α inhibitor (30 μM), AP-1 inhibitor (10 μM) or 30 min with ZAK inhibitor [3h (10 μM) and 6p (10 μM)] or IKKβ inhibitor BAY11-7082 (5 μM). The pretreated or untreated BMMs or A549 cells were transfected with active trimer 2-5A (2’,5’-p_3_A_3_), inactive dephosphorylated A3 (2’,5’-A_3_) ^53^ or with poly(I):poly(C) (Millipore Sigma, #528906) at the concentrations indicated in figure legends with Lipofectamine 2000 (Invivogen, #190772) or Lipofectamine 3000 (Invitrogen L3000001), according to the manufacturer’s instructions. THP-1 cells were transfected with 2-5A in reverse transfection manner adding cells to the lipofectamine/2-5A mixture according to the manufacturer’s instruction, and then seeding cells in 12-well plates. At the indicated time after transfection, cells were harvested either with RIPA buffer for Western blots or with total RNA Miniprep Super Kit for total RNA purification (Cat: B610583, Bio Basic) following the manufacturer’s instructions. Anisomycin (5 μM) treatments were done on THP-1 or A549 cells for 1h, and cells were harvested with RIPA buffer for Western blots.

### rRNA cleavage assays for RNase L activity

After transfection of 2-5A or pIC at the indicated concentrations and times, cells were lysed with RLT buffer and the total RNA was isolated with EZ-10 Spin Columns Total RNA Minipreps Super Kit. RNA was separated on RNA chips with an Agilent Bioanalyzer 2000 and the separation images were obtained.

### Bulk RNA-sequencing (RNAseq)

The WT and *Rnasel*^-/-^ BMM were mock-transfected or transfected with either 2-5A (2’-5’p_3_A_3_) (20 μM) or as a negative control A_3_ (2’-5’A_3_) (20 μM) in RNase-free water in the presence of lipofectamine and incubated for 3 h at 37°C in an incubator with 5% CO_2_. RNA was extracted, quantified in a uv spectrophotometer, and evaluated for integrity by RNA chip on an Agilent Bioanalyzer, model 2100. mRNA-Seq libraries were prepared from poly(A) tailed RNA, quantified and sequenced at Novogene. Primary analysis of the RNA-seq data was done at the Bioinformatic Core Facility, Florida Research & Innovation Center, Cleveland Clinic by Adrian Reich, and at the Institute for Computational Biology, School of Medicine, Case Western Reserve University by E. Ricky Chan. Briefly, the RNA-seq original sequence data FASTQ files were first inspected by fastqc to ensure data quality. The reads were trimmed of exogenous adapter sequences and low-quality bases were removed using cutadapt (version 2.8). The trimmed reads were aligned to the mouse genome (ENSEMBL GRCm38). Quantification of reads was performed to generate the gene-level feature counts from the read alignment with STAR (version 2.7.5a). The feature counts were further normalized for the gene size and library to obtain the gene expression value for all genes and all samples. The differential gene expression analyses were conducted by contrasting the 2-5A- or A_3_-transfected samples, with the mock-transfected samples by DESeq2 (version 1.30.1) and DEXSeq (version 1.36.0) using R (version 4.0.4). After obtaining the sequence reads and DESeq2 comparison tables (Supplementary Tables S2-S8), the gene expression data were plotted using volcano plot and heatmap by R program (genes list in heatmaps are shown in Supplementary Tables S10-S14), and the clustering of genes was based on complete linkage and the Euclidean distances of gene expression values. Gene set enrichment analysis (GSEA) was conducted by projecting the fold-change ranking onto Hallmark gene sets (Supplementary Table S9) (https://www.gsea-msigdb.org/gsea/msigdb/human/genesets.jsp?collection=H) and C5 ontology gene sets (https://www.gsea-msigdb.org/gsea/msigdb/genesets.jsp?collection=C5).

### qRT-PCR

Total RNA was extracted from BMM, THP-1 cells and A549 cells as described above. RNA was reverse transcribed with high-capacity cDNA reverse transcriptase kit (4368814, ThermoFisher). qRT-PCR was performed with oligonucleotide primers (Table S1) and Bullseye EvaGreen qPCR 2x MasterMix-ROX (BEQPCR-R, MidSci) using a BioRad qPCR system (CFX Opus 384). Results were analyzed by the ΔCt or ΔΔCt method, as described in the manufacturer’s instructions.

### ZAKα gel shift assays

A549 cells were pretreated with ZAK inhibitor compounds 3h (10 μM) or 6p (10 μM) for 30 min. The treated or untreated cells were then transfected with 2-5A (20 μM) for 2 h or treated with anisomycin (5 μM) for 1 h, and total protein was harvested with RIPA buffer. Proteins in cell lysates were separated on 7.5% Phos-Tag SDS-PAGE (Fujifilm Wako Chemicals U.SA. Corp, #AAL-107) following the manufacture’s instruction. The gels were then treated with 1 mM EDTA and the protein was transferred to PVDF membrane for Western blotting with anti-ZAKα antibody.

### Cell survival assays by real-time imaging of cells

Cells (2 × 10^4^ per well) were seeded in 24 well plates. After 18 h cells were mock transfected or transfected with 2-5A (20 μM) and stained for total cells with Syto^TM^ 60 red fluorescent nucleic acid stain (250 nM) (ThermoFisher Scientific, #S11342) (converted to blue by IncuCyte software) and for dead cells with Sytox^TM^ green nucleic acid stain (250 nM) (ThermoFisher Scientific, #S7020) in culture medium, then cells were cultured in an IncuCyte live–cell imaging system (SX5) for 60 h in an incubator at 37°C/5% CO2. The cell viability was calculated by counting the number of green objects per well (dead cells) which was then normalized to the total number of cells per well (red objects) and the averages of 9-16 images were presented as the final data.

## ACKNOWLEDGEMENTS

We thank E. Ricky Chan (Case Western Reserve University) and Adrian Reich (Florida Research and Innovation Center, Cleveland Clinic) for primary analysis of RNAseq data; Xiaoyun Lu (Jinan University, Guangzhou) for the generous gift of ZAK inhibitors, George Stark (Cleveland Clinic) and Bill Merrick (Case Western Reserve University) for discussions, Susan R. Weiss (University of Pennsylvania) for gifts of CRISPR constructs and cell lines, Babal K. Jha (Cleveland Clinic) for the mouse anti-RNase L antibody, and David Schumick (Enterprise Creative Services, Cleveland Clinic) for expert artwork. This research was supported by the National Institute of Allergy and Infectious Diseases of the National Institutes of Health under Awards R01AI135922 (to R.H.S.) and R01 AI104887 (to R.H.S and Susan R. Weiss) and to the Center for Gene Expression (CGEN) a Center of Excellence funded by The National Danish Research Foundation (grant no. DNRF166).

## LIST OF TABLES S2-S14 (Excel Files)

Table S1: PCR Primer Sequences.

Table S2: Raw transcript data from all analyzed transcriptomes.

Table S3: Normalized transcript data from all analyzed transcriptomes.

Table S4: Differentially expressed genes from all analyzed transcriptomes.

Table S5: Gene expression changes in transcriptomes from 2-5A transfected WT BMM vs mock-transfected WT BMMs.

Table S6: Gene expression changes in transcriptomes from A3-transfected WT BMM vs mock-transfected WT BMM.

Table S7: Gene expression changes in transcriptomes from 2-5A transfected RL KO BMM vs mock-transfected RL KO BMM.

Table S8: Gene expression changes in transcriptomes of A3-transfected RL KO BMM vs mock-transfected RL KO BMM.

Table S9: GSEA analysis of Hallmark gene sets with 2-5A transfected vs mock-transfected WT BMM.

Table S10: Interferon stimulated genes and antiviral genes heatmap transcript data

Table S11: Antigen presentation and process genes heatmap transcript data

Table S12: Inflammation genes heatmap data

Table S13: Ribosome protein genes heatmap data

Table S14: Housekeeping genes heatmap data

## SUPPLEMENTARY FIGURE LEGENDS

**Figure S1.**
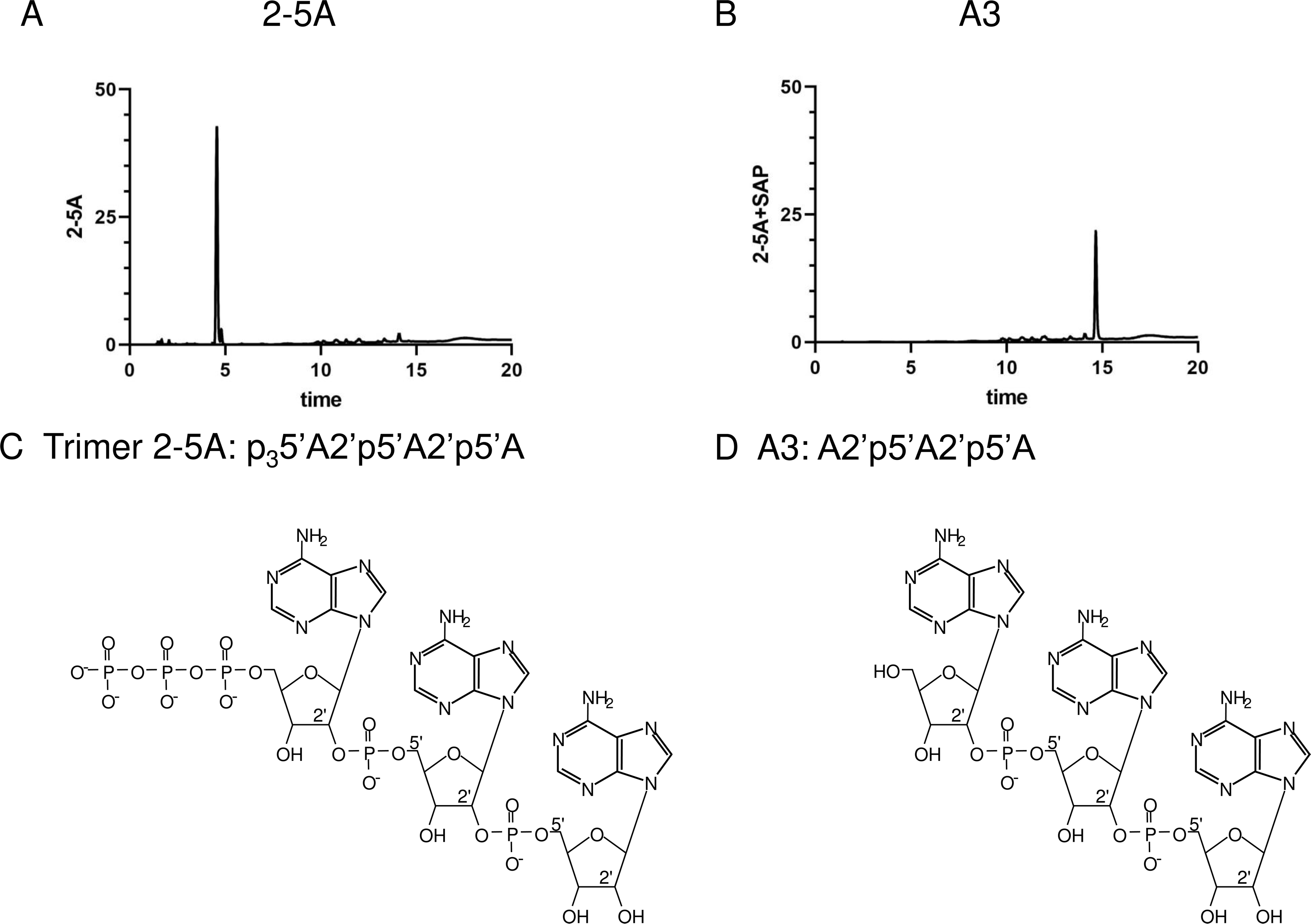
(A,B) HPLC analyses and (C,D) chemical structures of 2-5A and A3. (A-B) 2-5A and A3 purity analysis by HPLC. (C-D) Chemical structures of 2-5A and A3. SAP, shrimp alkaline phosphatase.

**Figure S2.**
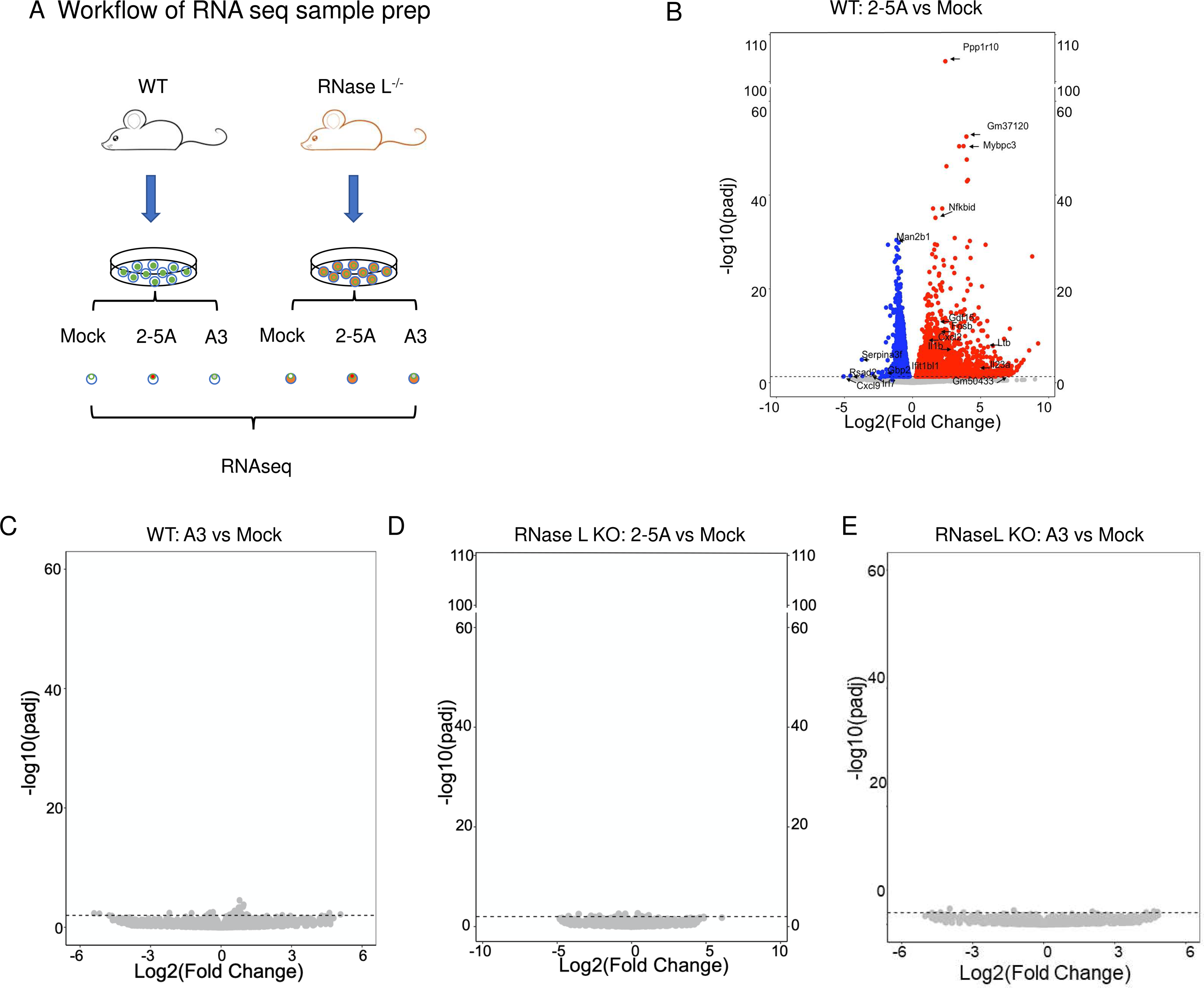
RNAseq experiments and data analyses from transfected WT and RNase L KO BMM. (A) Workflow diagrams for RNAseq experiments on WT and RNase L KO BMM that were mock-transfected, A3-transfected (20 μM, 3 h) or 2-5A-transfected (20 μM, 3 h). (B-E) Volcano plot of the transcript expression changes in comparisons of the different treatment groups.

**Figure S3.**
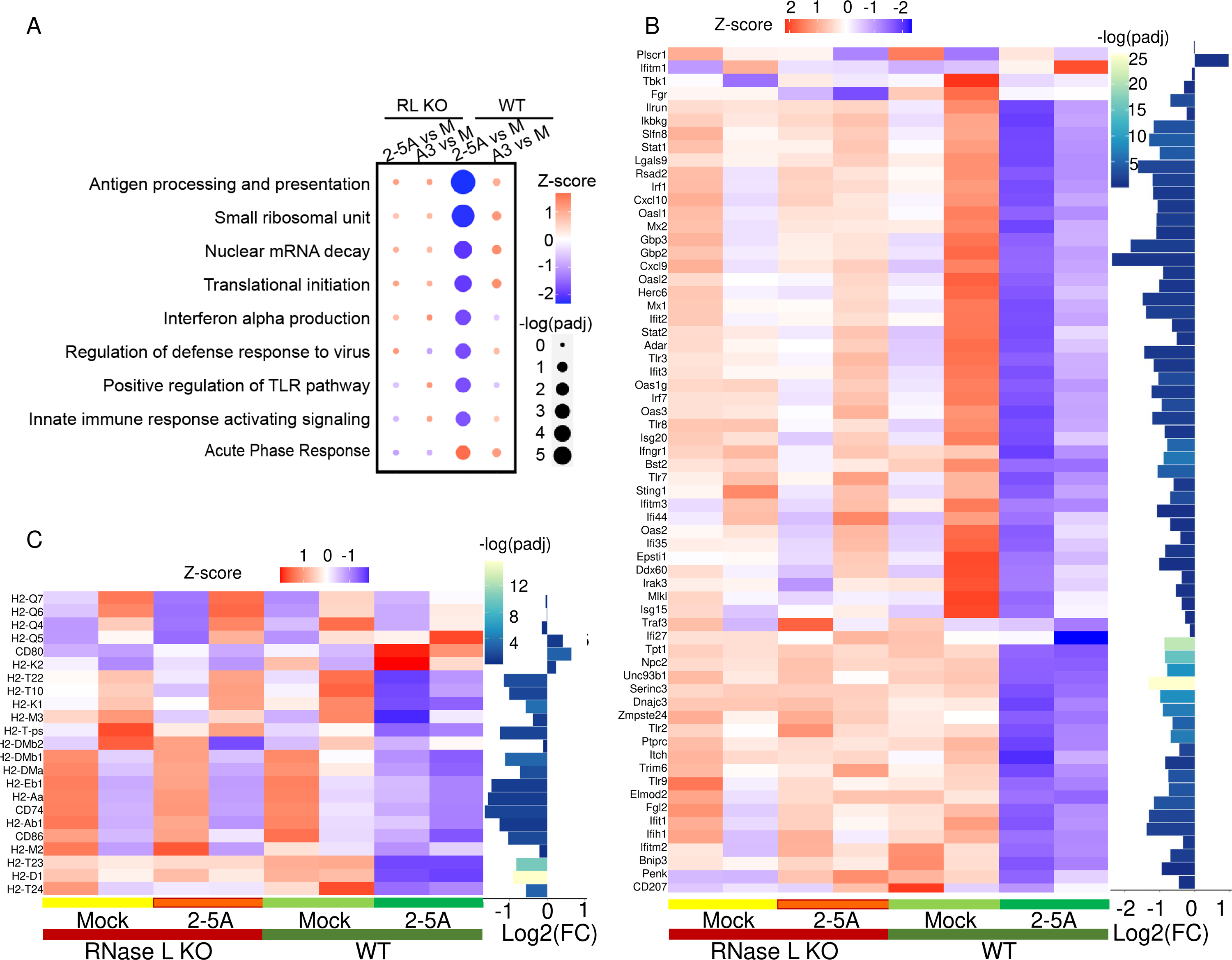
Regulation of transcript levels for antiviral genes, ISGs and antigen processing and presentation genes by 2-5A-transfection. (A) Gene ontology (GO) GSEA analysis of down-regulated and up-regulated transcript sets. (B) Heatmaps of transcripts for antiviral genes and ISGs and (C) antigen processing and presentation genes from BMM that were mock-transfected, or transfected with 2-5A or A3 (20 μM, 3h). Insert shows log2(fold change).

**Figure S4.**
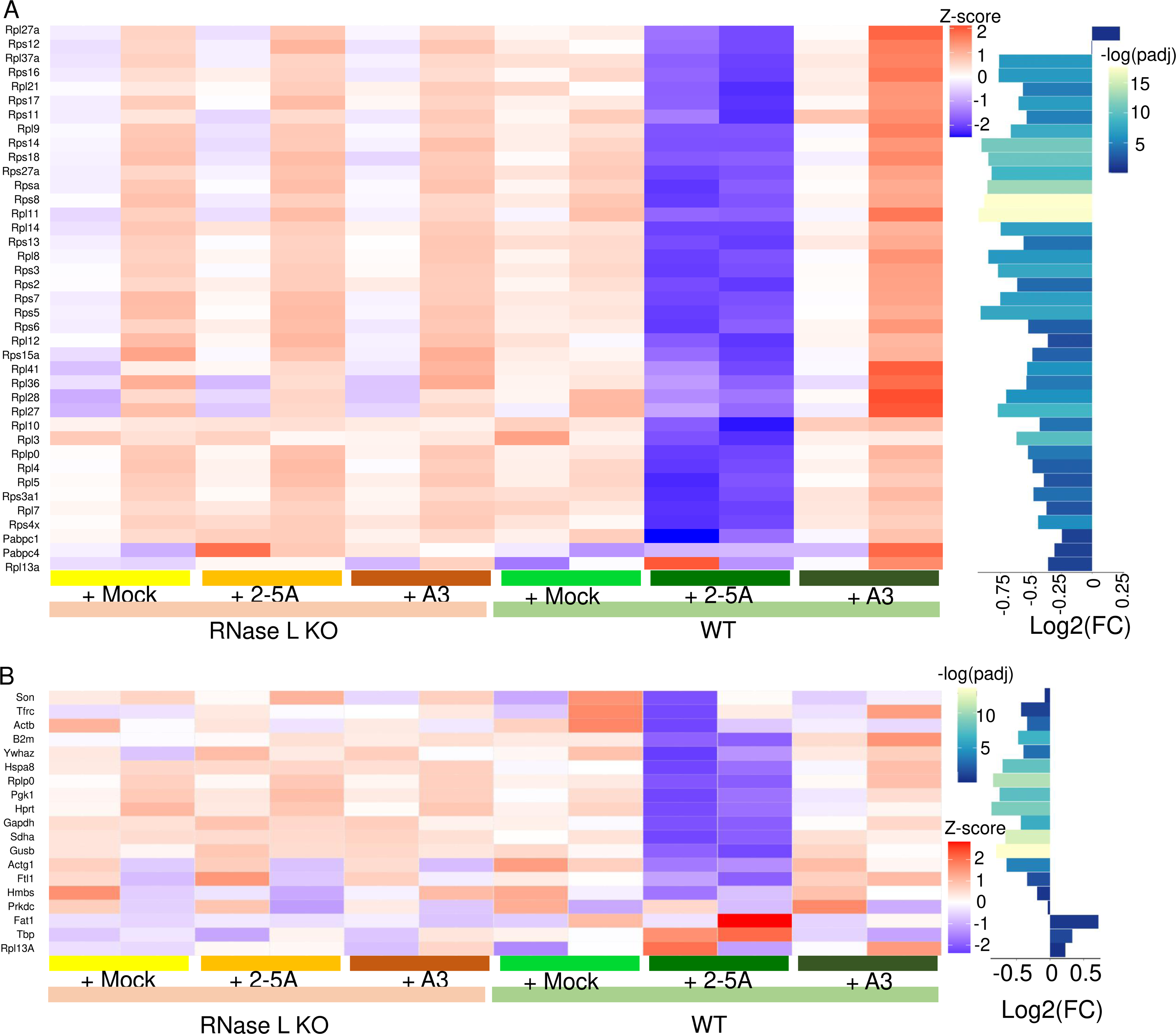
Regulation of transcript levels for ribosomal proteins and housekeeping genes by 2-5A-transfection. Heatmaps of transcripts for (A) ribosomal proteins, and (B) housekeeping genes. Inserts show z-scores and log2(fold change).

**Figure S5.**
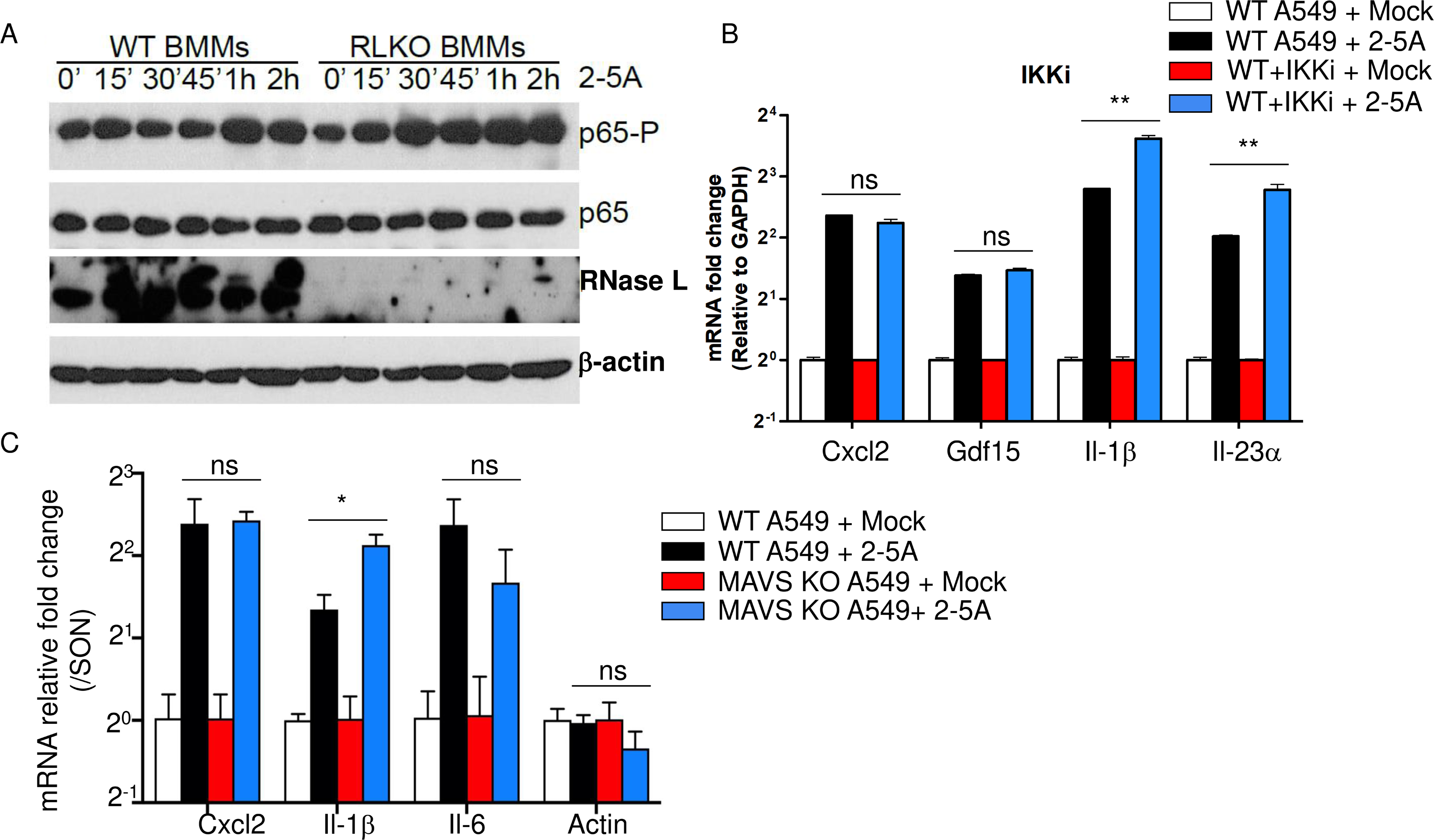
Lack of evidence for NF-κB and MAVS signaling during 2-5A induction of transcripts in A549 cells. (A) Levels of total and phosphorylated levels of p65 (subunit of NF-κB) after 2-5A transfections (20 μM) of WT BMM or RNase L KO BMM as determined in Western blots. (B) Relative levels of different transcripts (as indicated) by qRT-PCR normalized with GAPDH transcripts in WT A549 in the presence or absence of an inhibitor of IKKβ (IKKi) (5 μM added to cells 1 h prior to 2-5A transfections (20 μM, 3 h)). (C). Relative levels of different transcripts (as indicated) by qRT-PCR normalized to SON transcript levels in WT or MAVS KO A549 cells that were mock-transfected of transfected with 2-5A (20 μM, 3 h). RNase L KO, RL KO

**Figure S6.**
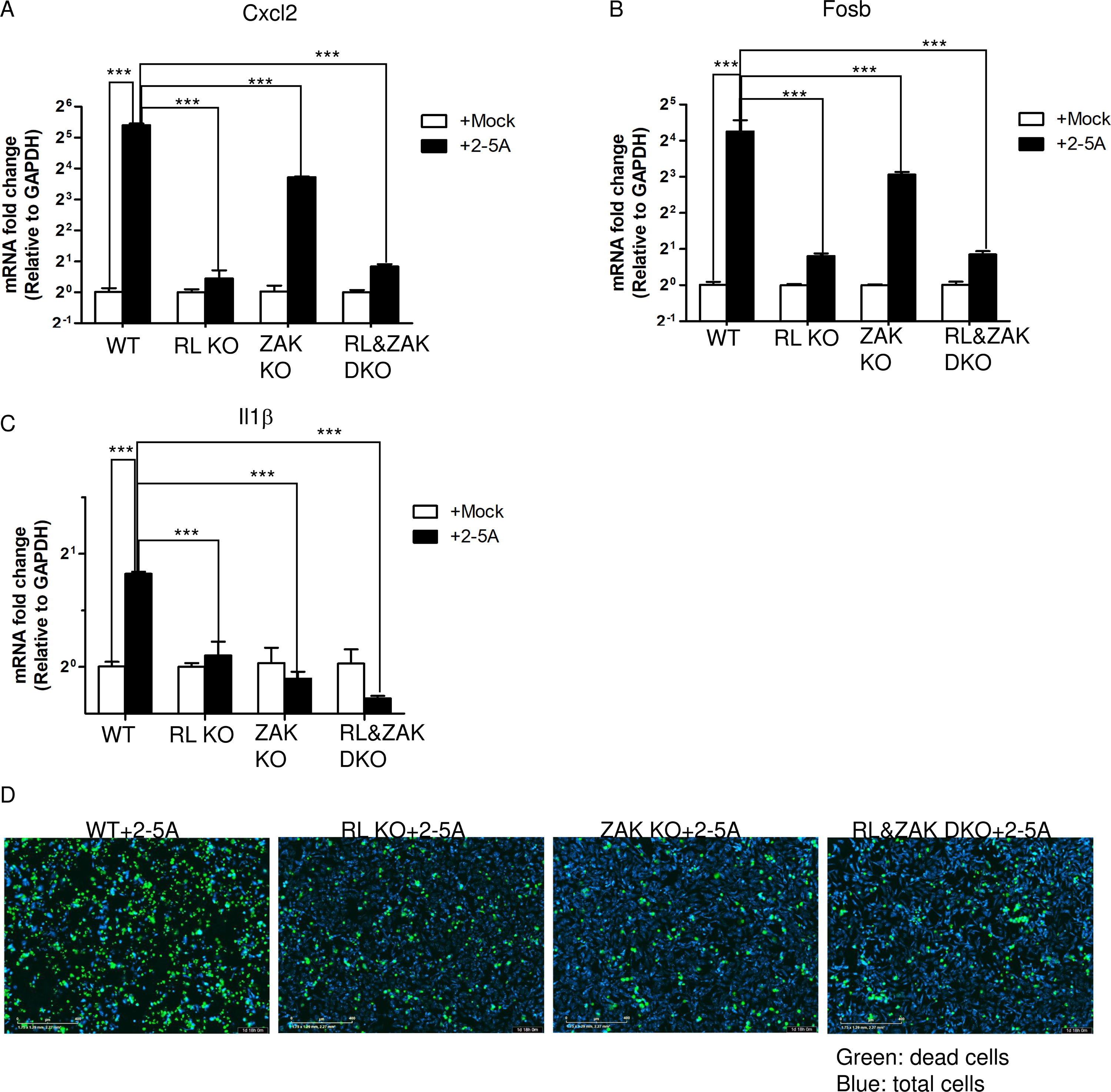
Regulation of transcript levels in WT, RL KO, ZAK KO and RL&ZAK DKO A549 cells after 2-5A transfections. (A-C). Relative levels of (A) Cxcl2, (B) Fosb, and (C) IL-1β transcripts normalized to GAPDH transcript levels after mock-transfections or transfection with 2-5A (20 μM, 3 h) of WT, RL KO, ZAK KO and RL KO/ZAK DKO A549 cells as determined by qRT-PCR. RNase L KO, RL KO; ZAK KO, ZKO; R&Z KO, RNase L/ZAK DKO. The experiment was repeated two times, each time in triplicate. Significance was determined by unpaired Student t-test. ***, *p*<0.001. (D) Representative images of stained A549 cell lines (as indicated) 42 h after mock-transfection or transfection with 2-5A (20 μM). Dead cells, green; total cells, blue.

**Table S1.**
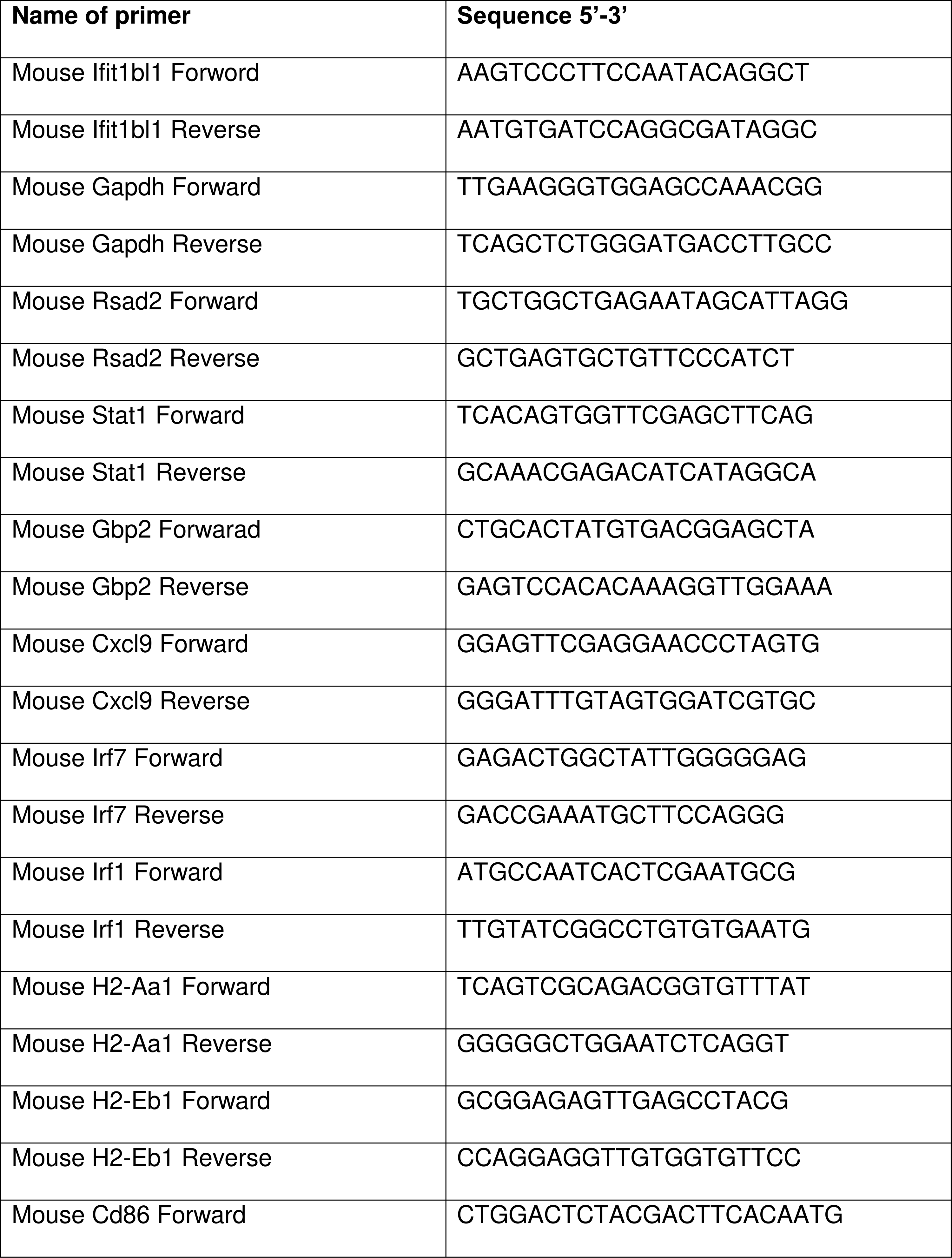
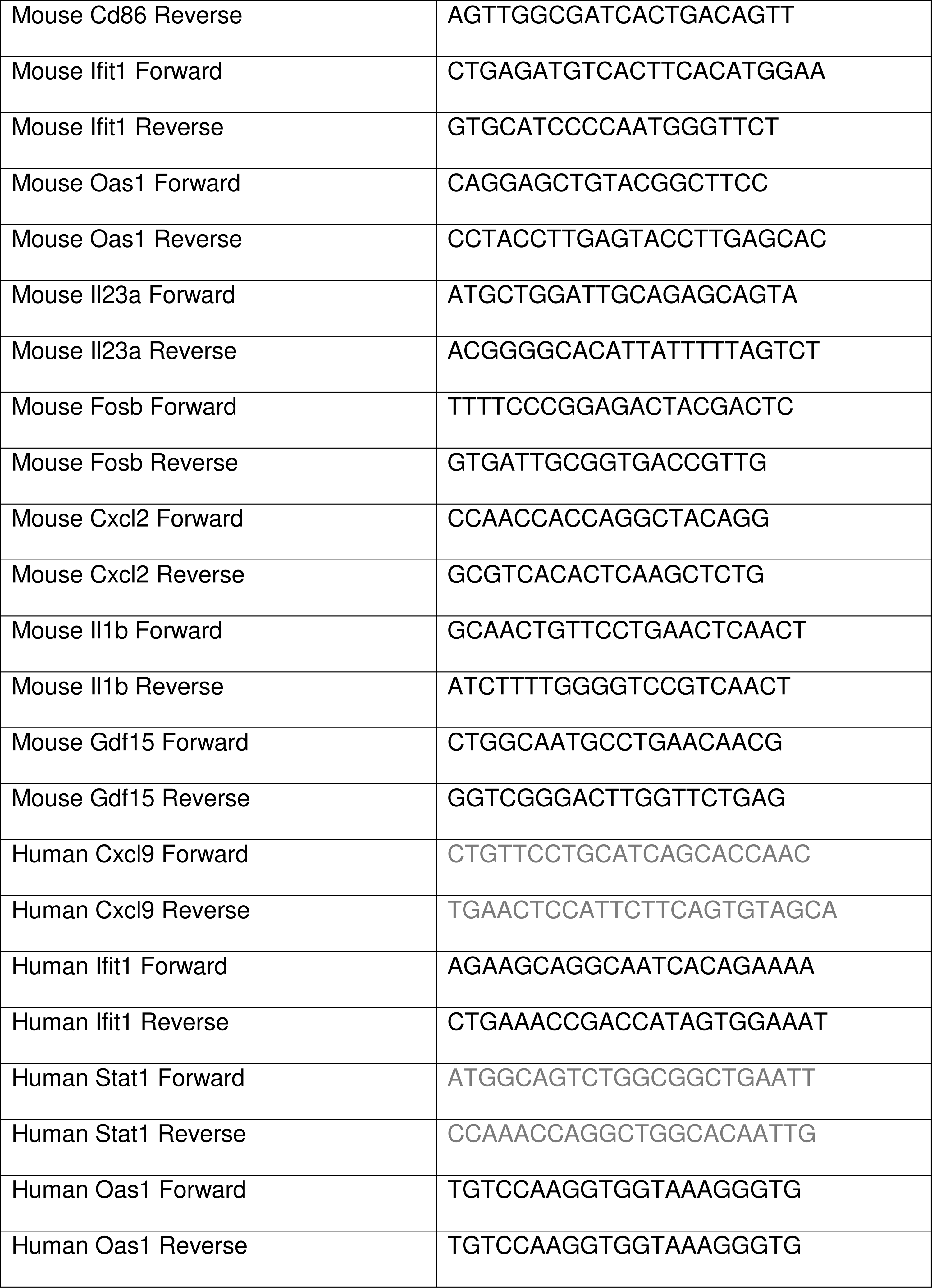
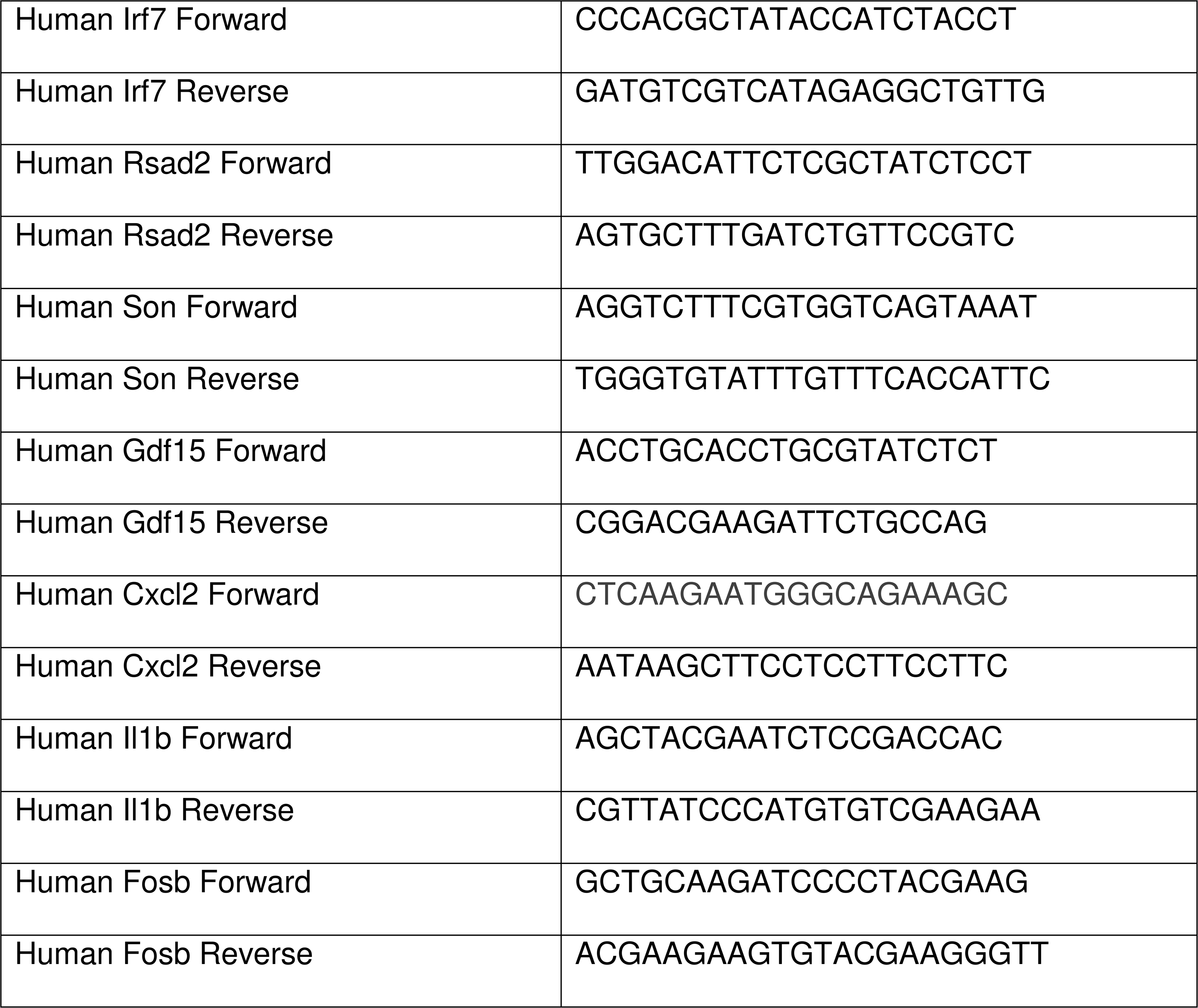
PCR Primer Sequences.

